# A unified ensemble-allosteric framework reconciles gain- and loss-of-function disease mutations in the IP_3_ receptor

**DOI:** 10.64898/2026.05.22.727127

**Authors:** Yu Zhu, Taufiq Rahman

## Abstract

Missense mutations in multidomain signalling proteins often produce divergent functional phenotypes despite preserved structural integrity, posing a fundamental challenge for interpreting human genetic variation. This problem is exemplified by the type 1 inositol 1,4,5-trisphosphate receptor, IP_3_R1, the principal neuronal IP_3_-gated Ca^2+^ release channel and a recurrent locus of pathogenic variation in *ITPR1*-associated ataxias and neurodevelopmental disorders. Although disease mutations cluster within the N-terminal suppression domain (SD) and IP_3_-binding core (IBC), how neighbouring substitutions can drive either gain- or loss-of-function remains unresolved. Here, we establish a unified ensemble-allosteric framework for IP_3_R1 channelopathy by integrating mutational constraint analysis, ensemble structural modelling, adaptive molecular dynamics, Markov state modelling and dynamical network analysis. We show that pathogenic variants preferentially occupy a stability-preserving regime, perturbing local sequence-structure compatibility without inducing global structural collapse. Thus, disease arises not from loss of fold, but from corruption of the conformational and allosteric logic that links ligand recognition to channel gating. The IP_3_-binding pocket forms a discrete multistate ensemble whose populations, physicochemical properties and kinetic connectivity are selectively remodelled by mutation. The loss-of-function variant R269W disrupts a cationic IP_3_-coordinating site, enriches non-permissive binding-pocket states and diverts conformational exchange through indirect kinetic routes. By contrast, the gain-of-function variant R36C preserves local pocket competence but weakens suppressor-domain restraint by rerouting long-range communication through extended, less efficient allosteric pathways. These findings reconcile opposing disease phenotypes within a single mechanistic model, showing that pathogenic variation can be encoded in altered ensemble probabilities and information flow rather than in static structural lesions.

## Introduction

Calcium is a universal and versatile intracellular messenger that regulates a diverse array of cellular processes including muscle contraction, secretion, metabolism, transcription, excitability, synaptic plasticity and cell fate(*1*). Among the major pathways that generate intracellular Ca^2+^ signals are the inositol 1,4,5-trisphosphate (IP_3_) receptors (IP_3_Rs) that are mainly expressed within the membrane of the endoplasmic reticulum (ER). Binding of the second messenger, IP_3_ and low (nanomolar) Ca^2+^ activates the IP_3_Rs, which then mobilise Ca^2+^ from the ER lumen into cytosolic Ca^2+^ transients with precise spatial and temporal profiles(*2*). IP_3_Rs are now understood not simply as passive conduits for stored Ca^2+^, but as signal-integrating hubs, whose activity is governed by binding of cytosolic ligands (IP_3_, Ca^2+^, ATP), interacting proteins and long-range allosteric coupling within the tetrameric channel complex(*2–4*).

Among the three mammalian isoforms, IP_3_R1, encoded by *ITPR1*, is especially important in the nervous system and is highly enriched in cerebellar Purkinje cells, where precisely regulated Ca^2+^ release is essential for dendritic integration, synaptic plasticity and motor control. The physiological importance of this particular IP_3_R isoform is underscored by experimental and human genetic evidence: mice lacking IP_3_R1 develop severe ataxia and seizures(*5*), disruption of IP_3_R1 in adult Purkinje cells destabilises spine morphology and cerebellar circuitry(*6*), and pathogenic *ITPR1* variants underlie a spectrum of neurological disease in human. Haploinsufficiency underlies SCA15(*7*), missense variants cause congenital non-progressive ataxia/SCA29(*8, 9*), and monoallelic or biallelic pathogenic variants also cause Gillespie syndrome(*10, 11*). Large clinical series further show that pathogenic missense variants cluster non-randomly in defined functional regions of the protein, particularly the N-terminal IP_3_-binding region, and that the phenotype extends beyond classical congenital ataxia to include broader developmental presentations(*12, 13*).

Mechanistically, the N-terminal region of IP_3_R1 (will be designated as R1-NT from henceforth) is a particularly informative site for understanding disease mechanism because it integrates ligand recognition, suppressive regulation and allosteric signal initiation within a single structural module but also harbours most mutations to date. Structural and biochemical studies resolved this region into an N-terminal SD, comprising approximately residues 1-225, and is immediately followed by the IP_3_-binding core (IBC) forming the minimal ligand-binding region(*16, 17*). The IBC itself is organised into β- and α-subdomains that close around ligand, whereas the SD modulates IP_3_ sensitivity and supports regulatory protein interactions(*16*). This arrangement means that the N-terminus is not simply the place where IP_3_ docks. Rather, it is an intrinsically allosteric module in which ligand recognition, suppression and signal transmission are already physically integrated before one even reaches the regulatory and transmembrane portions of the receptor.

This residue-level architecture provides a natural framework for interpreting pathogenic missense variation. Classical mutagenesis and structural analyses converged on a chemically coherent IP_3_-binding pocket in which residues in the β-subdomain, particularly R265, T267, G268 and R269, and residues in the α-subdomain, particularly K508, R511, Y567, R568 and K569, make major contributions to ligand recognition and phosphate coordination(*17, 18*). Full-length ligand-bound structures of IP_3_R1 refined this view by showing how ligand engagement is coupled to local pocket contraction and broader rearrangements across the cytosolic assembly(*15*). Disease-associated variants in the N terminus map onto several mechanistic layers of this architecture. Substitutions near D34 and R36 lie in the SD, whereas variants around R241, T267, R269, A275, S277, K279 and A280 occupy the IBC-β arm of the binding region, and variants such as E512K, Y567C, R568G and N602D lie on the opposing IBC-α side of the pocket. Thus, pathogenic residues do not all report on a single structural mechanism: some are positioned to alter suppressive control, whereas others are positioned to perturb ligand recognition or pocket-to-channel coupling.

This separation between SD regulation and direct ligand-pocket disruption is already evident in the functional literature. The SD variant R36C increases IP_3_-binding affinity and produces a gain-of-function (GOF) phenotype with altered intracellular Ca^2+^ signals, demonstrating that pathogenicity in R1-NT is not restricted to loss of channel activity(*19*). By contrast, systematic analysis of SCA29-associated substitutions within or near the N-terminal binding region showed that variants including R241K, T267M, T267R, R269G, R269W, S277I, K279E and A280D abolish channel activity, most commonly by impairing IP_3_ binding, whereas other N-terminal substitutions can preserve ligand binding yet disrupt downstream gating(*20*). The N-terminal mutation map therefore does not point to a single pathogenic mechanism; instead, it defines a mechanistically structured landscape in which neighbouring residues can differentially affect suppressive control, ligand recognition and signal propagation.

**Table 1.**
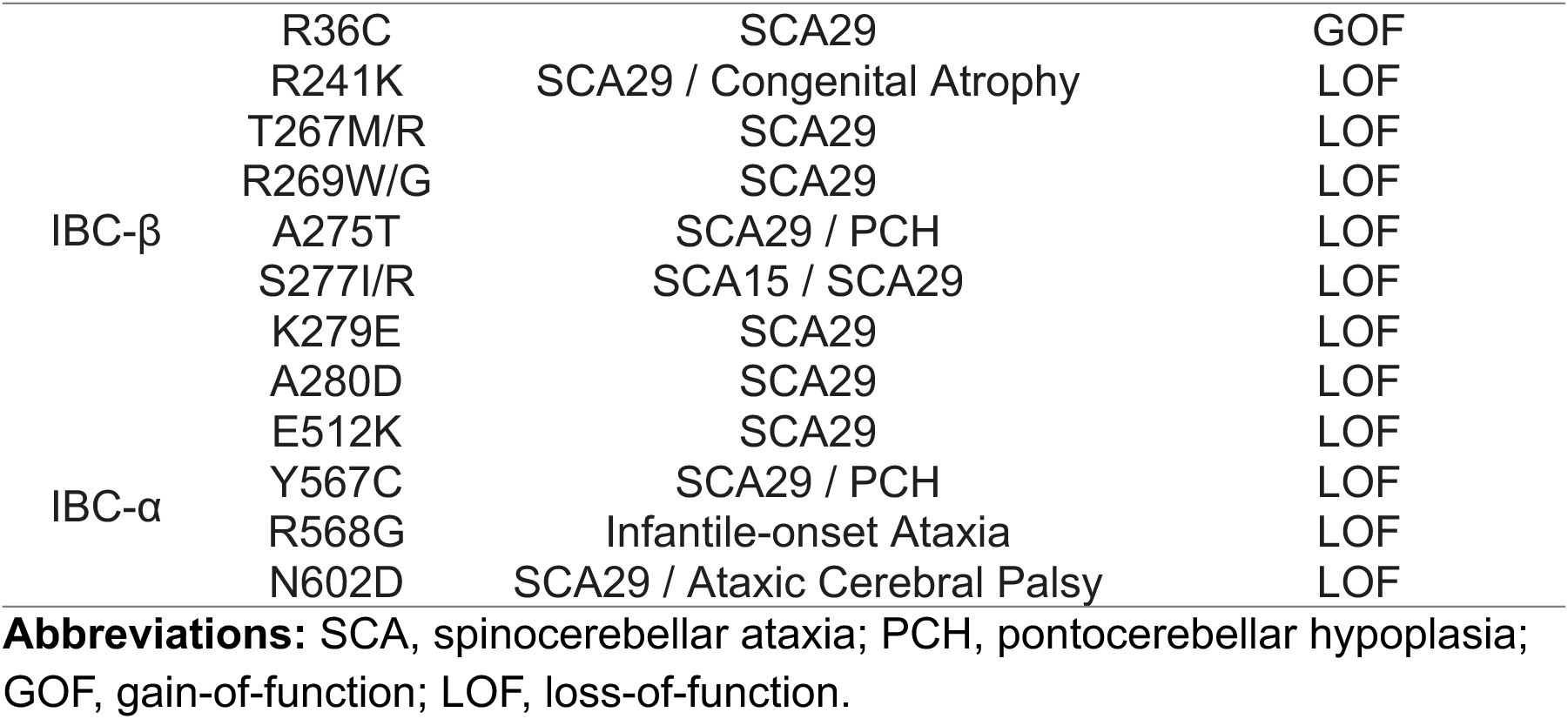
Reported disease-associated mutations in the R1-NT. Disease-associated variants are grouped by localisation within the SD or the β- and α-subdomains of the IBC (IBC-β and IBC-α) and annotated by reported disease association and functional consequence. Adapted from *Arige et al. 2025* with additional recent clinical reports(*13*).

Full-length structure studies have made this mechanistic distinction even more important. The cryo-EM reconstructions established the architecture of the tetrameric channel and revealed that the N-terminal domains of one subunit communicate extensively with neighbouring subunits through the cytosolic assembly(*21*). Ligand-bound structures then demonstrated that agonist engagement drives coordinated rearrangements within the IBC and across the broader allosteric scaffold, rather than simply occupying a static cleft. Complementary structures of human IP_3_R3 in multiple gating conformations, together with structural titration across a range of Ca^2+^ concentrations, further supported a conformational landscape view of IP_3_R activation gating in which the receptor samples resting, preactivated, activated and inhibited states(*15, 22*). Functional work further proved that productive Ca^2+^ release requires occupancy of all four ligand-binding sites within the tetramer(*23*). Taken together, these findings argue that IP_3_R function is encoded not in a single static structure but in the relative accessibility, connectivity and cooperative coupling of multiple conformational states.

This framework exposes a central gap in current understanding of *ITPR1* channelopathies. We now know where many pathogenic residues lie, which structural states the receptor can adopt, and that the variant effects depend on both mutation location and subunit context. What remains still far less clear is how individual missense substitutions in the N-terminal region reshape the dynamic regulatory landscape that links residue identity, local pocket geometry, kinetic accessibility and long-range allosteric communication. This problem is especially acute for mutations that preserve the overall fold of the N-terminus yet still produce profound functional consequences. Variants centred on D34 or R36 are well positioned to alter SD coupling without directly deforming the ligand pocket, whereas variants centred on T267, R269, Y567 or R568 are positioned to perturb ligand recognition or favour non-productive pocket conformations. A static structural description is therefore insufficient to explain how neighbouring substitutions can preserve the N-terminal fold while producing divergent GOF and LOFoutcomes. Instead, the relevant mechanistic question is how these mutations redistribute conformational ensembles, alter the kinetics of exchange between substates, and rewire communication between the SD and the IBC.

Here, we address this problem by integrating saturation mutagenesis, generative ensemble modelling, adaptive molecular dynamics, Markov state modelling and contact-informed dynamical network analysis to determine how disease-associated mutations reshape the regulatory landscape of the human R1-NT. Our premise is that pathogenicity in IP_3_R1 is not adequately explained by global destabilisation or by static structural inspection alone. Rather, it must be understood within a stability-constrained allosteric landscape in which amino-acid substitutions can selectively alter sequence-structure compatibility, reweight pocket substates, remodel kinetic connectivity and redistribute long-range communication between the SD and the IBC. Framed in this way, the R1-NT becomes a tractable system in which mutational biophysics, conformational regulation and disease mechanism can be analysed within a single coherent structural language.

## Results and Discussion

### A biophysical constraint landscape links sequence compatibility, stability, and pathogenicity in IP3R1 NT

To establish a biophysical framework for interpreting disease-associated mutations in the R1-NT (residues 1-604), we performed a comprehensive *in silico* saturation mutagenesis analysis across the SD and the IBC. For each possible single-amino-acid substitution, we quantified changes in sequence-structure compatibility using inverse folding likelihoods (ΔlogP) derived from ProteinMPNN(*24*), and changes in thermodynamic stability (ΔΔG) predicted using ThermoMPNN(*25*). In this framework, more negative ΔlogP values denote substitutions that are less compatible with the local folded environment, whereas more positive ΔΔG values indicate greater predicted destabilisation of the folded state. Together, these metrics define whether a mutation is likely to perturb the local structural code, the global energetic integrity of the fold, or both.

Across all substitutions, ΔlogP and ΔΔG exhibit a moderate-to-strong anticorrelation, with Spearman ρ ranging from −0.56 to −0.76 across method combinations (Fig. 2A and Fig. S1), consistent with prior evidence that protein sequence evolution is constrained by folding stability and mutational fitness(*26–28*). This relationship indicates that energetically favourable residues are not selected independently of structural context, but instead emerge from a coupled constraint space that simultaneously encodes fold stability and sequence compatibility. However, the resulting mutational landscape is continuous and highly structured, with no natural separation at ΔlogP = 0 or ΔΔG = 0. This absence of intrinsic thresholds argues against simplistic binary interpretations of mutational effects and instead supports a graded constraint landscape shaped by the architecture of the R1-NT. This indicates that mutational effects are governed by intrinsic features of the R1-NT rather than by simple energetic thresholds.

**Figure 1.**
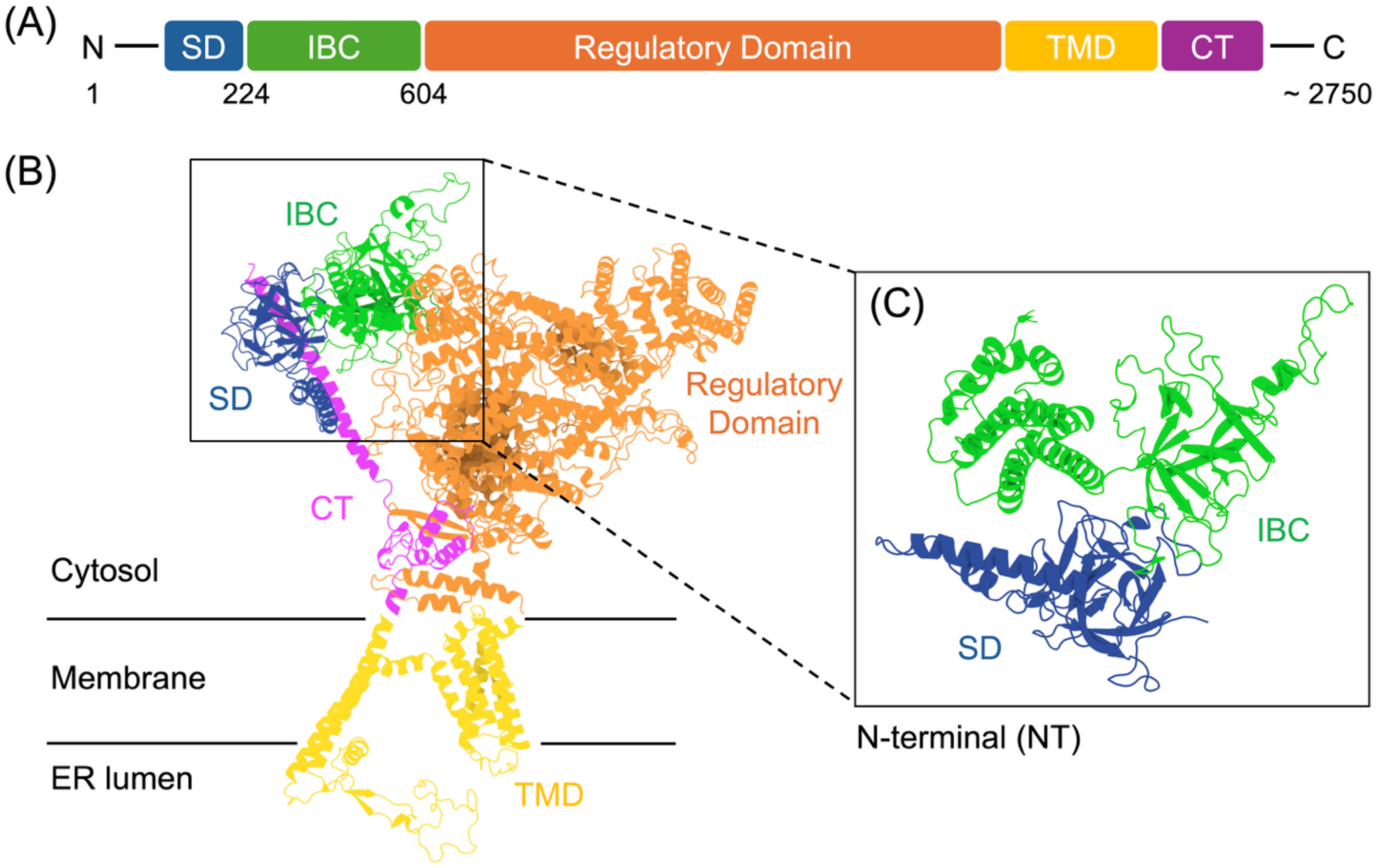
Domain organisation and cryo-EM architecture of the human IP_3_R1 monomer. **(A)** Linear schematic of the human IP_3_R1 sequence (UniProt Q14643). Functional domains are colour-coded: N-terminal suppression domain (SD, blue, residues 1-225); IP_3_-binding core (IBC, green, residues 226-604); regulatory domain (orange); transmembrane domain (TMD, yellow); and C-terminal tail (CT, purple). **(B)** Cartoon representation of the IP_3_R1 monomeric unit (extracted from the tetramer, PDB: 8EAR) showing its orientation relative to the ER membrane. The extensive cytosolic assembly, comprising the SD, IBC, and regulatory domains, communicates with the pore-forming TMD. Horizontal lines indicate the approximate boundaries of the phospholipid bilayer. **(C)** Zoomed-in view of the N-terminal (NT) region. The SD (blue) and IBC (green) are physically integrated to form a discrete allosteric unit. The SD sits at the cytosolic apex, where it is positioned to modulate ligand access to the IBC and facilitate long-range communication with the regulatory assembly. Molecular graphics were rendered in ChimeraX(*14*) from PDB 8EAR(*15*).

**Figure 2.**
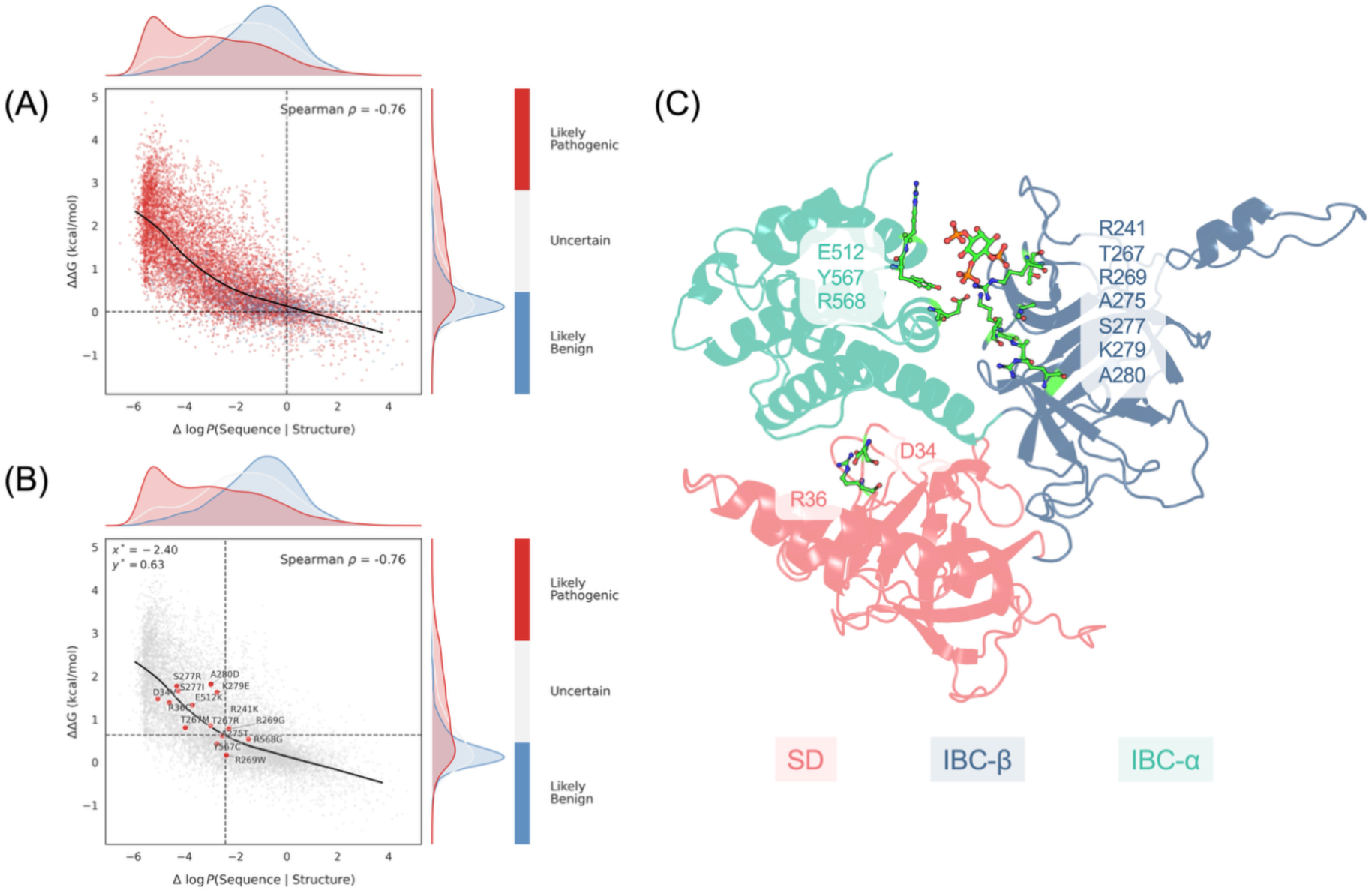
Saturated in silico mutagenesis and structural mapping of the R1-NT. **(A)** Mutational scan of the NT showing the correlation between sequence-structure compatibility and predicted thermodynamic stability. Points are coloured by the predictions from AlphaMissense: likely pathogenic (red), uncertain (grey), and likely benign (blue). Spearman’s ρ = −0.76. **(B)** Mapping of known gain- and LOF variants within a reparameterised coordinate system. Dashed lines indicate data-driven reference points (x* = −2.40, y* = 0.63) where pathogenic and benign distributions intersect. Most functional variants occupy a regime of reduced sequence-structure compatibility but modest destabilisation, preserving structural integrity. **(C)** Structural localisation of high-impact variants on the NT regulatory module, comprising the SD (salmon), IBC-β (blue), and IBC-α (green). Residues highlighted in (B) are shown as sticks, demonstrating their clustering at critical domain interfaces.

To relate this biophysical landscape to functional relevance, we next incorporated pathogenicity predictions from AlphaMissense(*29*) for all possible substitutions. The distribution of predicted pathogenicity was strongly skewed, with a large fraction of mutations predicted to be pathogenic and comparatively few classified as benign, consistent with the essential regulatory role of the R1-NT(*4, 16, 17, 30*). When the predicted pathogenicity scores are mapped onto the ΔlogP-ΔΔG landscape, pathogenic mutations are preferentially enriched in regions characterised by reduced sequence-structure compatibility, whereas benign mutations cluster toward regions of higher compatibility and lower destabilisation (Fig. 2A). Importantly, this enrichment was distributed across the joint ΔlogP-ΔΔG landscape rather than being separable by a single threshold on either axis, indicating that pathogenicity is encoded in relative positioning within the constraint space rather than absolute energetic penalties.

Since AlphaMissense-predicted pathogenic and benign substitutions did not segregate along the nominal zero axes (ΔlogP = 0 or ΔΔG = 0), we implemented a data-driven reparameterisation, translating the coordinate origin to the intersection (x*, y*) of their respective score distributions. This transformed coordinate system defines four mutational regimes relative to protein-specific reference values, preserving the physical interpretation of both ΔlogP and ΔΔG while capturing the intrinsic constraint structure of the R1-NT. Within this framework, the majority of pathogenic mutations occupy a regime characterised by strongly reduced sequence-structure compatibility but only modest destabilisation. In contrast, mutations predicted to be both highly destabilising and sequence-incompatible are comparatively rare. This observed asymmetry supports a model in which many pathogenic substitutions in the R1-NT occupy a stability-preserving incompatibility regime: they perturb local residue-environment compatibility while maintaining the overall folded architecture. Such mutations are therefore well positioned to disrupt ligand recognition or allosteric regulation without causing global structural collapse.

Mapping reported disease-associated *ITPR1* mutations onto the reparameterised landscape reveals strong concordance with this constrained regime (Fig. 2B). Both GOF and LOFvariants cluster in regions of low ΔlogP and moderate ΔΔG, consistent with functional disruption arising in the absence of severe destabilisation. Notably, the IP_3_-binding residue R269W occupies a distinct position characterised by low ΔlogP and low ΔΔG, highlighting it as a mutation that preserves thermodynamic stability despite substantial perturbation of local sequence compatibility. This positioning distinguishes R269W as a mutation that perturbs functional specificity without compromising structural integrity, suggesting a mechanism rooted in altered local interactions rather than global destabilisation. Structural mapping of high-impact variants onto the N-terminal regulatory module further supports this interpretation (Fig. 2C). Mutations cluster at interfaces between the SD and the IBC, as well as within the ligand-binding pocket itself. This spatial organisation reinforces the notion that pathogenic mutations target regions that mediate inter-domain communication and ligand recognition, rather than regions required for maintaining the global fold.

To assess the robustness of these conclusions, we repeated the analysis using alternative inverse folding, stability prediction, and pathogenicity scoring frameworks, including ESM-IF(*31*), FoldX(*32*), and ESM-1b(*33*) based metrics. Across all model combinations, the qualitative structure of the ΔlogP-ΔΔG landscape and the enrichment of pathogenic mutations within the stability-preserving regime were preserved (Fig. S2). This cross-model consistency demonstrates that the observed constraint landscape is not an artefact of any single predictive framework, but instead reflects an intrinsic property of the R1-NT.

Collectively, these analyses identify the R1-NT as a stability-constrained mutational landscape in which sequence-structure compatibility and thermodynamic stability are coupled but non-equivalent. Predicted and reported disease-associated substitutions converge on a regime of reduced local compatibility with modest destabilisation, arguing against global unfolding as the dominant mechanism of pathogenicity. Instead, these variants appear to act within an intact N-terminal scaffold, where local residue-environment incompatibility can alter ligand recognition, pocket-state probabilities or SD-IBC regulatory coupling. This reframes pathogenicity as a problem of fold-preserving conformational and allosteric dysregulation, motivating the ensemble-based, kinetic and network analyses that follow.

### Disease-associated mutations reweight IP3-binding pocket conformational ensembles within the retained native-like conformational manifold

To determine how disease-associated mutations perturb the conformational landscape underlying IP_3_ recognition, we generated ensemble representations of the R1-NT using BioEmu(*34*). For each system including wild type (WT) and selected IP_3_-binding-related variants within the NT, 1,000 conformations were sampled under identical conditions. Conformations exhibiting pronounced disruption of inter-subdomain contacts were excluded using a contact-based filtering scheme, thereby enriching for structurally coherent states that preserve native domain architecture while retaining local conformational variability. This filtering strategy ensured that subsequent comparisons were performed within a physically meaningful conformational manifold, minimising artefacts arising from non-native opening or model hallucination.

Given that IP_3_ recognition is mainly governed by subtle rearrangements within the ligand-binding pocket rather than large-scale domain motions(*15, 17, 35*), subsequent analyses focused on pocket-local structural features. This choice was guided by structural and functional evidence that the IBC behaves as a clam-shell-like recognition module, in which ligand engagement reshapes the relative arrangement of pocket-forming β- and α-subdomains before propagating to downstream gating transitions(*17, 23*). Each retained conformation was therefore represented as a reduced feature set comprising pocket-local backbone geometry, backbone dihedral angles, and side-chain 𝜒_1_ rotamers of predefined IP_3_ coordinating residues. Projection of all conformations into a low-dimensional embedding using UMAP(*36*) revealed a reproducible organisation of the pocket conformational space into discrete, densely populated regions (Fig. 3A), while variant-resolved distributions are shown in Fig. S3. A density-derived pseudo-free-energy reconstruction of the same embedding resolved three principal basins (Fig. 3B), indicating that the IP_3_-binding pocket samples a structured multi-basin landscape rather than a continuously diffusive ensemble. This observation accords with recent cryo-EM analyses showing that IP_3_Rs sample multiple preactivated and activated conformations, and that ligand binding biases the occupancy of pre-existing states rather than imposing a single deterministic structural transition(*15, 23, 35*).

**Figure 3.**
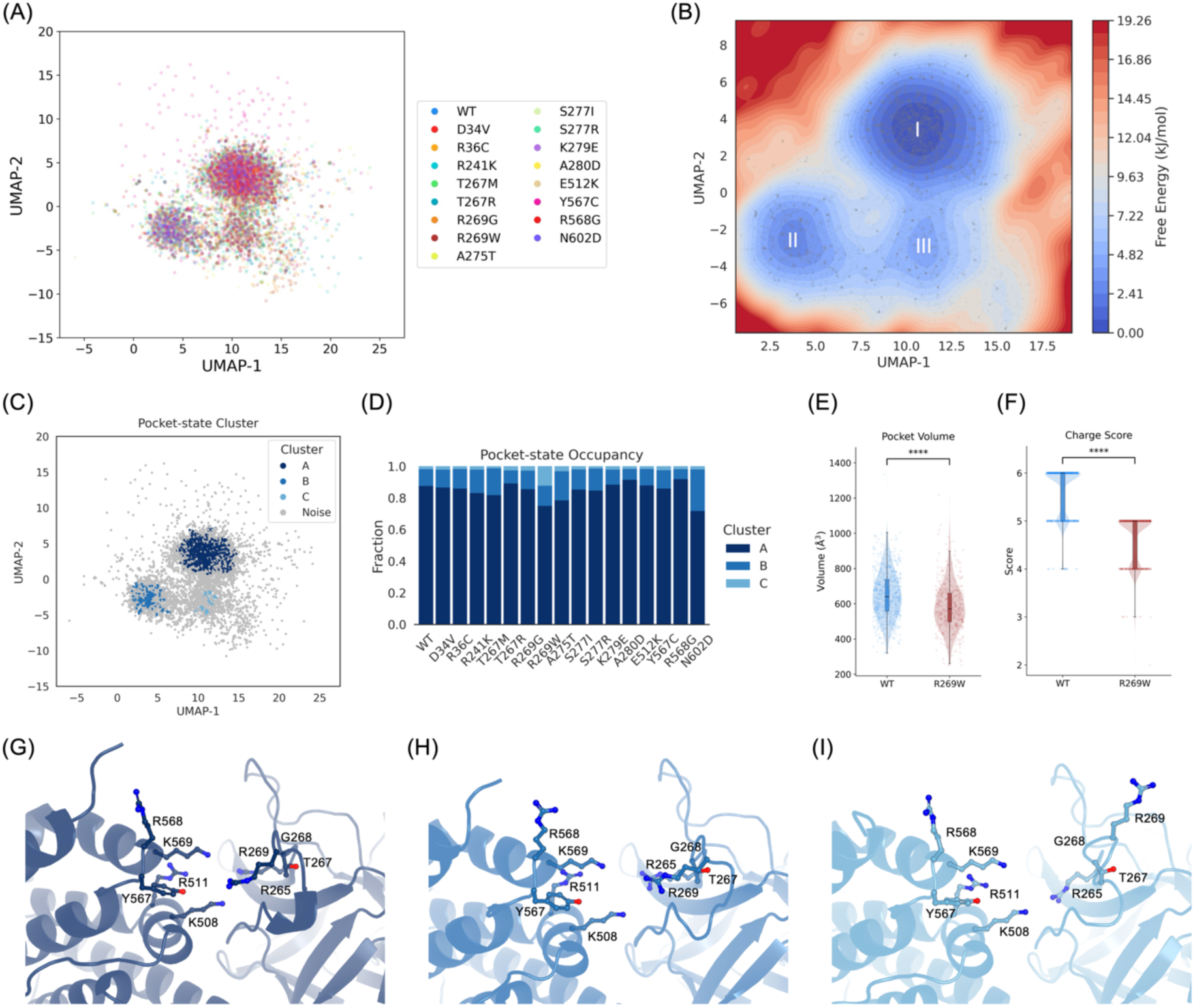
Disease-associated mutations reshape the conformational and physicochemical landscape of the IP_3_-binding pocket. **(A)** UMAP projection of BioEmu-derived pocket conformations for WT and disease-associated variants. **(B)** Pseudo free-energy surface of the combined ensemble, revealing three dominant conformational basins (I, II and III). **(C)** Density-based (DBSCAN) clustering of the same embedding identifies three geometrically defined pocket substates, labelled A-C. **(D)** Fractional occupancy of each substate across WT and variants. **(E)** Distribution of pocket volume in WT and R269W. **(F)** Distribution of pocket charge score in WT and R269W. **(G-I)** Representative structural snapshots of the three pocket substates, highlighting rearrangements of residues surrounding the IP_3_-binding site.

To obtain a data-driven partition of pocket conformational space, DBSCAN clustering was applied to the reduced pocket-local embedding (Figure 3C). DBSCAN was used because it does not require the number of clusters to be specified a priori and permits low-density regions to be treated as noise(*37*). Across a range of clustering parameters, three dominant clusters were consistently recovered after noise removal. We therefore refer to these geometrically defined groups as pocket substates A, B and C. Although the labels should not be interpreted as discrete thermodynamic states in the strict kinetic sense, their agreement with the density-derived pseudo-free-energy basins supports their use as reproducible descriptors of local pocket geometry for comparing variant ensembles.

We next quantified mutation-induced redistribution of the pocket ensemble by computing the fractional occupancy of each substate for all variants and comparing these values to the WT ensemble (Fig. 3D; Table S1). The latter was dominated by substate A, which accounts for 87.5% of sampled conformations, whereas substates B and C were comparatively minor, accounting for 10.6% and 1.9%, respectively. The two GOF SD variants examined (D34V and R36C) exhibited pocket substate distributions closely resembling the WT, with only modest shifts in relative populations. This is intriguing given both residues reside in the SD rather than within the IBC itself. Previous experimental work showed that R36C increases IP_3_-binding affinity and produces an enhanced Ca^2+^ release phenotype(*19*), consistent with the longstanding idea that the SD tunes ligand sensitivity through interdomain coupling(*16, 30*) rather than by directly contributing to the pocket chemistry. The preservation of pocket-state occupancy in D34V and R36C therefore argues that GOF behaviour can arise without wholesale remodelling of the local binding landscape, and instead points towards an allosteric mechanism that perturbs suppressive control over an otherwise native-like pocket ensemble.

In contrast, the LOF variants displayed heterogeneous but systematic redistribution of pocket substate populations. Several variants exhibited enrichment or depletion of specific substates relative to WT. These changes indicate that LOF phenotype is likely to stem from a reweighting of pre-existing conformational states, rather than the emergence of entirely novel structural classes. This proposition aligns well with the broader IP_3_R mutation-relevant literature, in which pathogenic substitutions typically impair IP_3_ binding or the transduction of allosteric signals without affecting the global architecture of the protein(*4, 20, 38*). The ensemble view emerging here therefore complements prior biochemical and Ca^2+^ imaging studies by providing a structural rationale for how mutations can compromise function without causing global unfolding.

Among all variants analysed, R269W exhibited the most pronounced and distinctive redistribution of the pocket substates. Relative to WT, substate A was markedly depleted, falling from 0.875 to 0.749, whereas substate C increased more than sixfold, from 0.019 to 0.123 (Table S1). In the embedded conformational space, R269W preferentially sampled a region only sparsely populated by other variants (Fig. 3A), indicating stabilisation of a rare pocket configuration. This behaviour is consistent with available structural models indicating that R269 directly coordinates IP_3_(*15*). The substitution with tryptophan introduces both steric bulk and altered electrostatic properties at the binding interface. Such physical disruption reconciles our findings with previous reports that R269W diminishes IP_3_-binding affinity and exerts dominant-negative effects(*38*), concordant with the requirement for stoichiometric ligand occupancy to drive productive channel opening(*39*). Collectively, our conformational analysis demonstrates that this mutation does not merely attenuate binding affinity in a static sense, but fundamentally reconfigures the ensemble towards a geometry that is infrequently sampled by the WT receptor.

To define the structural basis of the R269W-associated shift more explicitly, we performed a frame-wise physicochemical analysis of the ligand-binding pocket using fpocket(*40*) derived descriptors (Fig. S4; Table S2). R269W produced a significant reduction in pocket volume, with the median volume decreasing by approximately 70 Å^3^, while total solvent-accessible surface area (SASA) remained largely unchanged. This pattern argues against gross pocket collapse and instead indicates selective remodelling of cavity shape and internal composition. The most striking changes involved the chemical character of the cavity. The charge score decreased by one unit, the hydrophobicity score increased strongly, apolar SASA increased, polar SASA decreased, and the proportion of polar atoms was reduced. Taken together, these descriptors indicate that R269W converts the pocket into a more hydrophobic and less positively tuned environment, precisely the opposite of what would be expected to favour productive engagement of IP_3_, a highly polar and multiply phosphorylated ligand(*17*). From a mechanistic perspective, the reduction in positive electrostatic character is especially compelling because R269 directly contributes to the local interaction field that accommodates the phosphate groups of IP_3_. Additional pocket descriptors reinforced this interpretation. R269W decreased the number of alpha spheres, increased alpha sphere density and the proportion of apolar alpha spheres, and modestly shifted centre-of-mass distance metrics, together indicating a cavity that is more compact, more internally hydrophobic, and more spatially reorganised without becoming globally occluded. Importantly, many of these features showed large effect sizes, with Cliff’s delta (δ) values approaching or reaching the extreme range for several descriptors, indicating that the observed differences are not merely statistically significant but structurally substantial within the sampled ensemble. By contrast, polarity score and flexibility remained unchanged, and total SASA did not reach significance (Fig. S4). These negative results are equally informative, because they indicate that the mutation selectively remodels pocket chemistry and internal packing rather than producing indiscriminate disorder.

This distinction is important in light of recent structural work showing that IP_3_R activation proceeds through coordinated movements of the β-trefoil and armadillo-repeat domains within the IBC and the wider cytosolic assembly(*15, 17, 23*). IP_3_ binding does not simply occupy a static cleft but it biases the receptor towards preactivated states that favour subsequent Ca^2+^-dependent gating. The present data suggest that R269W perturbs this process at an earlier stage by altering the local conformational and physicochemical profiles of the binding pocket itself. In this premise, this mutation appears to reduce the probability of sampling ligand-compatible microstates before any productive coupling to the broader activation pathway can occur. This interpretation aligns with the functional characterisation of pathogenic variants mutated within the IBC. Substitutions surrounding the binding pocket frequently manifest as LOF alleles, either by attenuating ligand affinity or by uncoupling ligand binding from downstream gating transitions.

More broadly, the ensemble-level effects observed here align with an emerging view of large signalling proteins in which function is encoded in the relative populations of interconverting conformational states rather than in a single privileged structure. For IP_3_Rs specifically, cryo-EM and functional studies now support a conformational landscape model in which IP_3_, Ca^2+^ and ATP bias the receptor across resting, preactivated, activated and inhibited ensembles(*23, 35*). Some pathogenic mutations can be mapped directly onto this framework. While GOF substitutions in the suppressor domain leave the local pocket landscape largely unaffected, LOF variants, particularly those at residues that directly coordinate IP_3_ or stabilise the IBC, shift the equilibrium towards non-permissive pocket states. These conformational shifts provide a physical basis for the disruption of downstream cellular processes. IP_3_R-mediated Ca^2+^ release regulates cellular excitability, secretion, mitochondrial metabolism, and triggers the store-operated Ca^2+^ entry(*4*), while mutations across all three IP_3_R subtypes are linked to neurological disorders, immunodeficiency, and exocrine dysfunction(*4, 41*). Within this context, the pocket remodelling induced by R269W describes how a single missense mutation in *ITPR1* disrupts intracellular Ca^2+^ signalling to result in the cerebellar phenotypes characteristic of SCA29. Conversely, the near-WT pocket distributions of D34V and R36C suggest that GOF phenotypes are driven primarily by altered SD communication. This hypothesis is explored in subsequent sections through explicit allosteric network analysis.

Together, these results establish that IP_3_R1 function is governed by the population balance among discrete pocket substates, rather than by preservation of a single static pocket structure. The two GOF SD variants examined here largely preserved the native pocket ensemble despite altered signalling output, whereas LOF mutations redistribute conformational weight among substates. In particular, R269W demonstrates a substantial ensemble reprogramming coupled to local physicochemical remodelling. These findings provide a mechanistic link between the biophysical constraint landscape and the functional defects observed in pathogenic variants. Rather than inducing catastrophic destabilisation, mutations within the IP_3_-binding region reshape the probabilities, geometries, and chemistries of pre-existing pocket states, thereby altering ligand compatibility and subsequent channel activation. This ensemble-centred perspective provides a coherent framework for interpreting both historical mutagenesis data and newly emerging clinical alleles. To determine whether these population shifts are accompanied by changes in the kinetics of interconversion between pocket substates, substate connectivity and transition rates were subsequently quantified using Markov state modelling.

### Disease-associated mutations remodel the kinetic connectivity of IP3-binding pocket substates

Having identified that disease-associated variants reweight IP_3_-binding pocket substates, we next asked whether these population shifts are accompanied by altered kinetic connectivity. We therefore performed adaptive MD simulations of the WT R1-NT, R269W and R36C. For each system, adaptive sampling generated 2 μs of aggregate simulation and was guided by pocket-local collective variables defined as explicit pairwise Cα-Cα distances among residues forming the ligand-binding pocket. This design enriched sampling of conformational exchange within the IP_3_-recognition site while preserving the surrounding N-terminal architecture. The kinetic analysis was restricted to a structurally predefined pocket formed by residues R265, T267-R269, K508, R511 and Y567-K569, thereby anchoring the model to the ligand-binding interface rather than to global domain displacement. This residue set includes R269, a central IP_3_-contacting residue whose substitution to tryptophan produced the strongest pocket-state redistribution in the ensemble analysis.

Because ensemble reweighting does not reveal whether pocket states remain kinetically accessible, we constructed Markov state models (MSMs) from the adaptive trajectories to quantify metastable pocket states, equilibrium populations, transition kinetics and dominant exchange routes(*42, 43*). MSMs provide a statistical framework for reducing high-dimensional MD trajectories into transitions among kinetically coherent conformational states, enabling estimation of state populations, mean first-passage times and transition pathways from simulation data. PyEMMA(*42*) was used for model estimation and kinetic analysis, including metastable-state assignment, Bayesian uncertainty estimation(*44*) and transition-path analysis. Model validation was performed using feature-set comparison, VAMP-2 scoring(*45*), implied-timescale analysis(*43*) and Chapman-Kolmogorov tests, supporting the stability of the inferred kinetic models across selected lag times and discretisation parameters (Fig. S5-S7). R269W was prioritised as the principal LOF case because it directly alters the ligand-binding interface(*15, 41*) and produced the strongest redistribution of BioEmu-derived pocket substates. R36C was included as a GOF comparator because it preserves a broadly WT-like pocket ensemble despite enhanced receptor activity, allowing local pocket kinetic disruption to be distinguished from distal SD-mediated regulation.

The WT MSM resolved three metastable pocket states arranged within a connected kinetic landscape (Fig. 4A-C; Table S3). The dominant state, S_3_, accounted for 48.3% of the equilibrium population, whereas S_1_ and S_2_ contributed 25.8% and 25.9%, respectively. Projection onto the leading independent components revealed three partially separated basins superposed on a continuous free-energy surface, consistent with a pocket that samples discrete conformational preferences without becoming kinetically fragmented (Fig. 4B). Transition-path decomposition was calculated from the least-populated state to the most populated state, providing a consistent comparison of how each system accesses its dominant basin. In WT, exchange from S_1_ to S_3_ was dominated by a direct route, accounting for 80.0% of transition flux, with only 20.0% proceeding through S_2_ (Fig. 4C; Table S3B). Pairwise MFPTs were in the tens to low hundreds of nanoseconds, indicating efficient interconversion among metastable pocket states (Fig. S5D; Table S3C). These results define a WT reference landscape in which the IP_3_-binding pocket maintains a preferred basin while retaining rapid access to alternative conformations.

**Figure 4.**
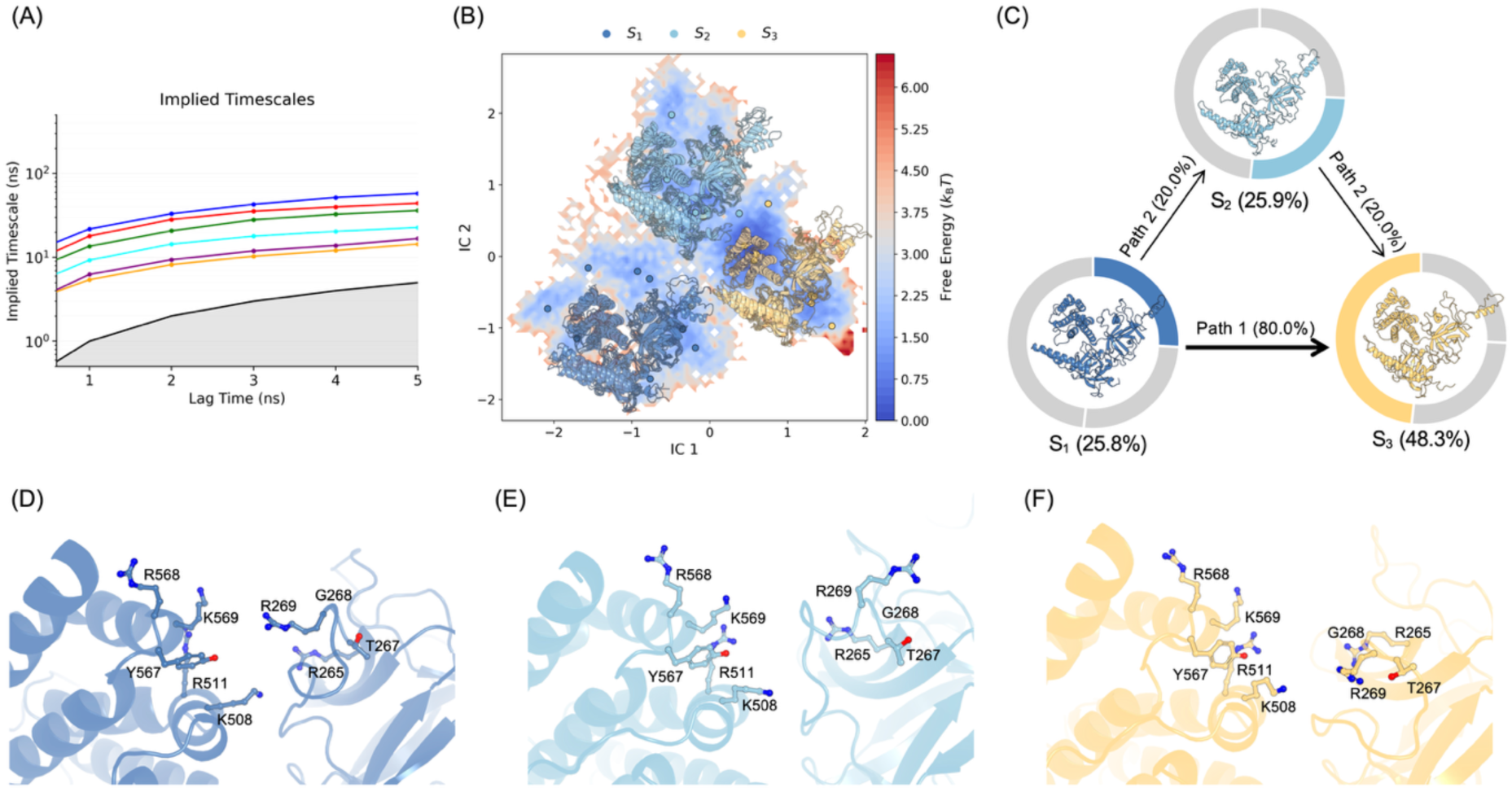
Conformational dynamics of the WT R1-NT reveal a three-state kinetic landscape within the IP_3_-binding pocket. **(A)** Implied timescale analysis of the WT Markov state model (MSM), validating kinetic separability across the selected lag time. **(B)** Projection of adaptive sampling trajectories onto the first two independent components (tICs), overlaid with the reconstructed free-energy surface and metastable state assignments (S_1_-S_3_). **(C)** Transition path decomposition from the least- to the most-populated state. Node areas reflect the equilibrium populations of S_1_-S_3_, with arrows indicating the dominant transition fluxes. **(D-F)** Representative conformations of states S_1_ (D), S_2_ (E), and S_3_ (F), focused on the IP_3_-binding pocket. Key ligand-coordinating side chains are shown as sticks and labelled.

Representative structures from the three WT metastable states provided a structural interpretation of this kinetic landscape (Fig. 4D-F). The predefined pocket residues adopted distinguishable arrangements across S_1_, S_2_ and S_3_, with changes in the relative positioning of R265, T267, G268, R269, K508, R511, Y567, R568 and K569. These structures suggest that the three kinetic basins correspond to expanded-like, intermediate-like and constricted-like pocket arrangements rather than globally distinct domain conformations. This interpretation is consistent with the preceding ensemble analysis, in which the same pocket region occupied a three-basin conformational landscape. The representative states therefore connect geometric pocket substates to kinetic exchange routes and provide a physical basis for the MSM topology.

R269W substantially altered this kinetic architecture (Fig. 5; Table S3). The mutant also resolved three metastable states, but their organisation differed from WT. The most populated state, S_3_, accounted for 41.3% of the ensemble, with S_1_ and S_2_ contributing 28.8% and 29.9%, respectively (Fig. 5C). Although this distribution indicates that R269W does not immobilise the pocket into a single static conformation, the free-energy surface and transition topology reveal a pronounced reorganisation of kinetic accessibility (Fig. 5B, C). In contrast to WT, where direct least-to-dominant exchange prevails, R269W shifted 86.8% of transition flux through an intermediate-mediated S1-to-S2-to-S3 route, leaving only 13.2% of flux on the direct path (Fig. 5C, H; Table S3B). This rerouting indicates that R269W reduces direct kinetic communication between pocket basins that are more readily connected in the native receptor.

**Figure 5.**
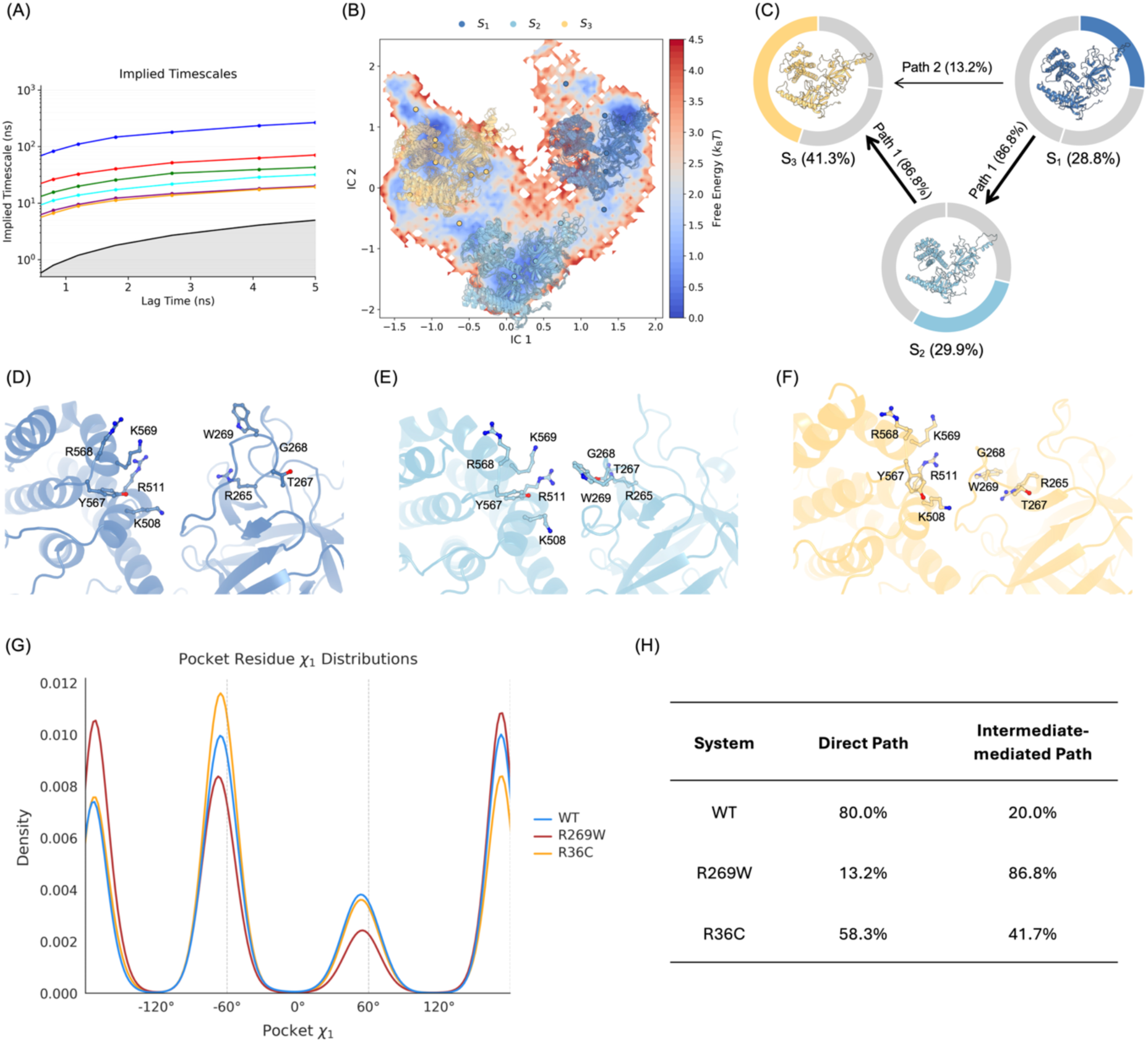
The R269W mutation reroutes pocket-state exchange and remodels the ligand-binding cavity. **(A)** Implied timescale analysis of the R269W Markov state model (MSM). **(B)** Projection of R269W adaptive sampling trajectories onto the first two independent components (tICs), overlaid with the reconstructed free-energy landscape and assigned metastable states. **(C)** Transition path decomposition from the least- to the most-populated state. Node areas indicate equilibrium populations, and arrows denote dominant transition fluxes, revealing a preference for an intermediate-mediated S_1_-S_2_-S_3_ route over direct S_1_-S_3_ exchange. **(D-F)** Representative conformations of the R269W metastable states, focusing on pocket residues. W269 and neighbouring residues are visualised as sticks. The dominant state exhibits repositioning of W269 and the adjacent loop, driving steric reshaping of the ligand-binding cavity. **(G)** Pooled 𝜒_!_distributions of pocket residues for the WT, R269W, and R36C variants. **(H)** Quantification of direct and intermediate-mediated transition fluxes from the least- to the most-populated state across the three variants.

The structural basis of this rerouting was apparent in the representative R269W metastable states (Fig. 5D-F). Replacement of the arginine with tryptophan introduces a bulky hydrophobic side chain at the centre of the ligand-recognition interface(*15*) and removes a positively charged group suited for engagement of the phosphorylated IP_3_ ligand. In the dominant R269W state, W269 and the adjacent T267-A275 loop adopt a conformation that encroaches upon the pocket, with neighbouring residues including T267, G268 and the region around A275 repositioned relative to the WT basin. Because A275 is itself a reported disease-associated residue(*4, 13*), involvement of this local segment suggests that mutation at R269 perturbs a broader pocket-regulatory loop rather than a single contact point. The resulting arrangement is consistent with steric reshaping, and potentially partial obstruction, of the ligand-binding cavity. Side-chain rotamer analysis further supported this interpretation (Fig. 5G). The pooled 𝜒_!_ distribution of pocket residues showed canonical rotameric wells in all systems, but R269W shifted their relative occupancy, with enrichment of trans-like 𝜒_!_conformations near ±180° and reduced sampling around the +60° well. By contrast, R36C retained a distribution closer to WT. Thus, the R269W effect extends beyond backbone-level pocket rearrangement to the side-chain organisation that defines the local steric and chemical environment of the ligand-binding site. This is particularly important for IP_3_ recognition, because productive binding depends on a constellation of polar and positively charged interactions within the binding core(*17*).

The kinetic effect of R269W is biologically consistent with prior functional studies. Comprehensive mutational analysis of disease-associated IP_3_R1 variants showed that R269W disrupts IP_3_ binding and abolishes IP_3_-induced Ca^2+^ release(*20*), supporting a severe loss-of-function mechanism. Additional disease-oriented studies and reviews distinguish ligand-binding variants from mutations that alter channel gating or inter-domain regulation, consistent with the separation between R269W and R36C observed here. The present simulations provide a dynamic explanation for these observations. R269W remodels the local pocket geometry, alters rotameric sampling and redirects state-to-state exchange through an intermediate route, thereby reducing the probability of direct transitions between ligand-relevant conformations. In this model, loss of function does not require global destabilisation of the N-terminal fold. Instead, R269W imposes a combined physicochemical, steric and kinetic penalty on productive IP_3_ recognition.

R36C displayed a distinct profile and was retained as a GOF comparator rather than as the central local-pocket phenotype (Fig. S7; Table S3). Its MSM also resolved three metastable states, with one dominant basin accounting for approximately half of the ensemble, but the exchange network remained more accessible than that of R269W. In the revised transition-path analysis, flux from the least-populated to the dominant R36C state remained weighted towards the direct route, but with a larger intermediate-mediated component than WT, 58.3% direct and 41.7% intermediate-mediated (Fig. S7C; Table S3B). This redistribution indicates that R36C subtly alters local pocket exchange, but without producing the strong intermediate-dominated rerouting, W269-driven steric remodelling or rotameric reorganisation observed for R269W. The local pocket landscape of R36C is therefore perturbed, but not locally disabled.

This distinction is consistent with the position and reported biology of R36C. R36 lies in the SD rather than the IBC, and R36C has been reported as a GOF mutation that increases IP_3_-binding affinity and alters Ca^2+^ signal patterns(*19*). The MSM analysis therefore suggests that R36C does not act by collapsing or occluding the IP_3_-binding pocket. Instead, its modest redistribution of local exchange, combined with preservation of broadly WT-like pocket structure, points to altered coupling between the SD and the IBC as the more plausible origin of enhanced activity.

This interpretation aligns with the broader structural view of IP_3_R activation. Cryo-EM studies(*15, 23, 35*) have shown that IP_3_Rs populate ligand- and Ca^2+^-dependent conformational ensembles, including resting, preactivated, activated and inhibited states, rather than a single rigid active structure. Within this landscape framework, a mutation such as R269W can impair channel activation by reducing access to ligand-compatible pocket states or by rerouting exchange through less favourable intermediates, even when the domain remains folded. Conversely, a SD mutation such as R36C can preserve local pocket kinetics yet alter the probability that ligand-binding events are transmitted to downstream regulatory motions.

The adaptive simulations and MSM analysis therefore establish two separable dynamical mechanisms. WT samples three connected pocket states with efficient interconversion. R269W redirects transitions through an intermediate-mediated route and stabilises a dominant pocket arrangement involving W269 and the adjacent T267-A275 loop, providing a local kinetic mechanism for loss of function. R36C preserves comparatively accessible pocket exchange but displays enough redistribution to suggest altered regulatory coupling rather than direct pocket failure. This distinction provides the rationale for moving from local pocket kinetics to contact-informed dynamical network analysis: if R36C does not primarily disable the ligand-binding pocket, its GOF effect is more likely encoded in the long-range communication pathways that couple the SD to the IBC.

### Gain-of-function mutation R36C weakens suppressive allosteric control by redistributing long-range communication through suboptimal pathways

R36C presents a mechanistic problem distinct from the pocket-centred LOF mutation R269W. The ensemble and MSM analyses showed that R269W remodels the IP_3_-binding cavity directly, whereas R36C retains a comparatively accessible, broadly WT-like local pocket landscape despite its GOF phenotype(*19*). This separation between local pocket competence and functional outcome implicates a regulatory mechanism operating through the SD, where residue 36 is located, rather than through direct obstruction of the ligand-binding site. Such a mechanism is consistent with experimental and disease-focused studies reporting that R36C increases IP_3_-binding affinity and alters Ca^2+^ release behaviour, whereas R269W disrupts IP_3_ binding and IP_3_-induced Ca^2+^ release(*20*).

Global patterns of correlated motion were first evaluated using dynamic cross-correlation matrices. This analysis provided a coarse-grained view of how each mutation alters correlated residue fluctuations across the N-terminal regulatory module. R269W produced extensive reorganisation of correlated motions across the SD and the IBC, consistent with its direct remodelling of the ligand-binding pocket and its severe LOF phenotype (Fig. S8A, B). In contrast, the R36C correlation pattern remained broadly similar to WT, with no evidence of widespread dynamical collapse or global rearrangement of the R1-NT (Fig. 6A, B). Thus, R36C is unlikely to act through global dynamic disorder of the N-terminal module. Instead, the preserved global correlation architecture points to a subtler mechanism in which specific SD-to-pocket communication routes are redistributed.

**Figure 6.**
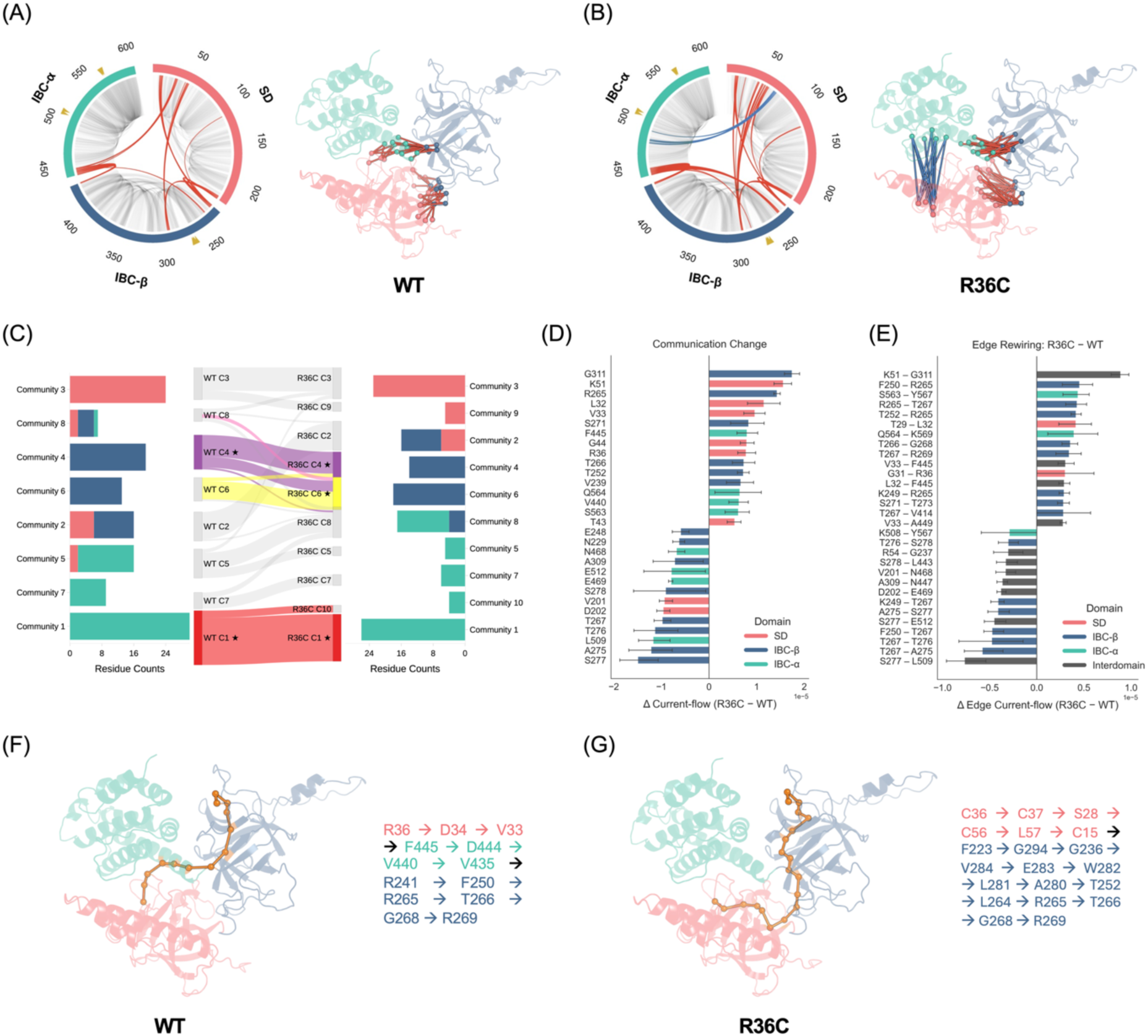
The R36C mutation preserves global correlated motions whilst redistributing SD-to-pocket communication via extended allosteric pathways. **(A)** Dynamic cross-correlation analysis of the WT R1-NT, presented as a domain-resolved circular representation and mapped onto the NT structure. The SD, IBC-β, and IBC-α are denoted in red, blue, and teal, respectively. Highlighted correlations delineate prominent coordinated motions characterising inter-domain and inter-subdomain interactions. The gold triangle designates the binding pocket residues. Corresponding dynamic cross-correlation analysis for the R36C mutant. The overarching correlation architecture remains broadly analogous to that of the WT, indicating that the R36C substitution does not induce global dynamic disruption. **(C)** Network community correspondence between the WT and R36C, derived from contact-informed dynamic networks. Pocket-associated communities are highlighted, demonstrating partial conservation alongside substantial reassignment of the residues facilitating source-to-pocket communication. **(D)** Residue-level alterations in current-flow betweenness centrality between R36C and the WT. Positive values denote increased source-to-pocket information flow in R36C, whereas negative values indicate attenuated flow. Error bars represent block-resolved variance. **(E)** Edge-level alterations in current flow, delineating inter-residue contacts that exhibit enhanced or diminished communication weights within the R36C variant. **(F)** The optimal weighted source-to-pocket communication pathway in the WT, mapped onto the NT structure and detailed by residue sequence. **(G)** The optimal weighted source-to-pocket pathway in the R36C mutant, illustrating an extended trajectory that encompasses additional SD and inter-domain residues prior to reaching the binding pocket.

To resolve this regulatory layer beyond global correlation maps, contact-informed dynamical networks were constructed from adaptive MD trajectories of the WT and R36C. The aim was to test whether a mutation like R36C that preserves the overall correlation architecture can nevertheless alter the efficiency, routing and modular organisation of communication between the SD and the IBC. In these networks, residues were represented as nodes, and edges were retained only when Cα-Cα contacts were persistent across the trajectory, thereby constraining communication routes to recurrent physical interactions. Each edge was weighted by the product of contact persistence and the magnitude of the corresponding dynamic cross-correlation coefficient, with edge length defined as the negative logarithm of this weight. This representation integrates structural proximity with correlated motion and avoids the overconnectivity of purely correlation-based networks. Current-flow betweenness was then used to quantify communication between a source region centred on residue 36 and target residues forming the IP_3_-binding pocket, because current-flow approaches integrate over ensembles of possible paths rather than relying on a single shortest route.

In the WT network, SD-to-pocket communication followed a compact route that bridged the SD, IBC-β and the IP_3_-binding pocket (Fig. 6A, F). Shortest-path analysis from R36 to individual pocket residues showed relatively direct access to both pocket lobes. For example, paths to R265, T267, G268 and R269 passed through D34, V33, F445, D444, V440, V435, R241 and F250 before entering the 265-269 pocket segment. Paths to K508, R511 and K569 followed a second compact route through D34, V33, F445, A446, Q513, E512 and R511. Across the evaluated pocket targets, the mean best weighted path length was 1.67, consistent with efficient native coupling between the suppressor domain and ligand-binding core (Table S7).

R36C increased the effective communication cost to every evaluated pocket target. The corresponding shortest paths from C36 were longer and rerouted through additional SD and inter-domain residues. Routes to the R265-R269 pocket segment passed through C37, S28, C56, L57, C15, F223, G294, G236, V284, E283, W282, L281, A280, T252 and L264 before reaching R265 and adjacent pocket residues. Routes to the K508/R511/K569 side of the pocket also shifted, using D34, V33, D448, N447, Q513, E512 and R511. The mean best path length increased from 1.67 in WT to 2.49 in R36C, a mean increase of 0.81 weighted length units, or approximately 1.5-fold (Table S7). Block-resolved analysis independently supported this conclusion, with R36C showing consistently longer best source-to-pocket paths across trajectory blocks (Fig. S8C). Thus, R36C does not simply replace the WT route with an equivalent alternative pathway but increases the cost of SD-to-pocket communication.

Node-level current-flow analysis identified the residues responsible for this redistribution (Fig. 6D; Table S5). R36C increased current flow through several SD and inter-domain residues, including G311, K51, L32, V33, G44, C36 and F445, as well as through pocket-proximal residues such as R265, S271, T266 and T252. In contrast, current flow decreased at residues within or adjacent to the ligand-binding pocket, including S277, A275, L509, T276, T267, S278 and E512. This pattern indicates redistribution of signalling responsibility rather than uniform strengthening or weakening. In particular, increased current through R265 and T266, coupled with reduced current through T267, A275, S277 and E512, suggests that R36C reroutes communication at the edge of the ligand-binding pocket rather than simply amplifying the native source-to-pocket pathway.

Edge-level current-flow changes provided a complementary view of this rewiring (Fig. 6E; Table S6). Several edges gained current flow in R36C, including K51-G311, F250-R265, S563-Y567, R265-T267, T252-R265, Q564-K569, T266-G268 and T267-R269. Conversely, edges such as S277-L509, T267-A275, T267-T276, F250-T267, S277-E512, A275-S277 and K249-T267 lost current flow. These altered contacts span the SD, inter-domain bridges, predefined pocket residues and immediately adjacent pocket-regulatory loops. More importantly, rewired edges involve the pocket residues R265, T267, G268, R269, Y567 and K569, as well as neighbouring disease-associated or regulatory positions such as A275, S277 and E512. This pattern indicates that R36C does not act through a single local contact but redistributes communication across the broader pocket-control network.

Community-level analysis provided structural context for the altered communication routes (Fig. 6C; Table S4). In WT, the predefined pocket residues were distributed across two principal communities: K508, R511, Y567, R568 and K569 clustered within an IBC-α pocket-side module, whereas R265, T267, G268 and R269 clustered within an IBC-β pocket-loop module. In R36C, the IBC-α pocket-side community was largely preserved, but the IBC-β pocket residues were redistributed: T267, G268 and R269 remained grouped, whereas R265 shifted into a separate pocket-adjacent community. This selective reassignment is notable because R265 lies at the entry of the R265-R269 ligand-binding segment and appears repeatedly in the altered source-to-pocket paths and edge-current changes. Thus, R36C does not dismantle the pocket communities wholesale, but changes how the two pocket lobes are modularly coupled to the SD. The highlighted Sankey analysis shows that pocket-involved communities are partially retained but their boundaries and inter-community relationships are reorganised, consistent with altered modular coupling between the SD and the IBC.

The best-path maps make this reorganisation physically interpretable. In WT, the source-to-pocket path runs through a relatively short chain from R36 to D34, V33 and the 443-445 region before entering IBC-β and the pocket-proximal 265-269 segment (Fig. 6F; Table S7). In R36C, the corresponding route is displaced toward a longer path involving C37, S28, C56, L57, C15 and a distal IBC-β arc spanning F223, G294, G236, V284, E283, W282, L281, A280, T252 and L264 before reaching R265, T266, G268 and R269 (Fig. 6G; Table S7). This detour provides a structural explanation for the increased weighted path length and supports a model in which R36C weakens direct SD control by forcing communication through less efficient inter-domain routes.

The comparison with R269W further clarifies the mechanistic distinction. R269W produced extensive network redistribution centred on the ligand-binding interface (Fig. S8A, B), consistent with its local pocket remodelling, altered rotameric sampling and LOF behaviour. R36C, by contrast, did not produce the R269W-like occlusive pocket phenotype. Instead, it preserved a broadly accessible pocket while increasing the communication cost between the SD and ligand-binding site. The two mutations therefore perturb different layers of the same regulatory architecture: R269W compromises the local ligand-recognition landscape, whereas R36C weakens the transmission of suppressive control from a distal regulatory domain.

This interpretation accords with structural models of IP_3_R activation in which ligand binding, Ca^2+^ and other regulatory inputs bias an ensemble of resting, preactivated, activated and inhibited conformations rather than switching the channel between two rigid states(*15, 23, 35*). Within such an ensemble-regulated system, gain of function need not arise from direct stabilisation of a ligand-bound pocket. Instead, weakening an inhibitory communication route from the suppressor domain can increase ligand sensitivity by reducing the effectiveness of suppressive coupling. R36C therefore provides a mechanistic example of gain of function through allosteric decoupling: the pocket remains locally competent, but the regulatory pathway that restrains it is redistributed through longer and less efficient routes.

These analyses complete a mechanistic separation between local and allosteric disease mechanisms in IP_3_R1 NT. R269W impairs ligand recognition by reshaping pocket geometry, chemistry and kinetic exchange. R36C acts through network-level reorganisation, increasing source-to-pocket path length, redistributing current-flow through compensatory residues and fragmenting pocket-associated communities. Rather than strengthening the ligand-binding pocket itself, R36C appears to weaken the native suppressive pathway that restrains pocket activation. This mechanism explains how a SD mutation can enhance IP_3_ sensitivity without producing the pocket-trapping signature observed for R269W, and links the biophysical constraint landscape, ensemble reweighting, MSM kinetics and allosteric communication into a unified model of mutation-driven IP_3_R1 dysregulation.

## Conclusions

Our study identifies the IP_3_R1 N-terminus as a stability-constrained allosteric space in which disease-associated missense variants perturb function not by collapsing the folding, but by selectively reweighting the conformational and communication states through which ligand recognition is coupled to channel regulation. Across mutational constraint mapping, ensemble modelling, adaptive MD, Markov state modelling and dynamical network analysis, pathogenic substitutions converged on a regime of reduced sequence-structure compatibility with preserved structural integrity, explaining how clinically severe *ITPR1* alleles can produce profound Ca^2+^-signalling defects without overt destabilisation of the N-terminal module. Within this framework, R269W and R36C exemplify mechanistically separable routes to disease. R269W acts locally, replacing a cationic IP_3_-coordinating residue with a bulky hydrophobic side chain, thereby remodelling pocket chemistry, enriching non-permissive binding substates and rerouting pocket exchange through less direct kinetic pathways. R36C acts distally, preserving a largely competent binding pocket while weakening suppressor-domain control by displacing native allosteric communication onto longer and less efficient routes. These findings reconcile GOF and LOF phenotypes within a single ensemble-allosteric model: pathogenic *ITPR1* mutations do not simply impair IP_3_R1 structure, but retune the probability, kinetics and directionality of regulatory information flow. This provides a mechanistic framework for interpreting emerging *ITPR1* variants and illustrates a broader principle by which missense mutations in multi-state signalling proteins convert preserved structural integrity into disease-relevant functional dysregulation.

## Materials and Methods

### Structural model and mutational landscape analysis

The N-terminal region of human IP_3_ receptor type 1 (IP_3_R1), encompassing the SD and the IBC (residues 1-604), was modelled using the reference structural model generated by BioEmu(*34*). This model was selected to ensure consistency between static structural analyses and subsequent ensemble-based conformational sampling. Side chains were reconstructed for all residues using standard rotamer libraries, followed by restrained energy minimisation to relieve local steric clashes while preserving the global fold. Residue numbering follows the canonical human IP_3_R1 sequence (UniProt(*46*) ID: Q14643).

Comprehensive single-amino-acid saturation mutagenesis was performed *in silico* across all positions in the modelled region. For each residue, all 19 non-WT substitutions were generated. Sequence-structure compatibility was quantified using inverse folding likelihoods computed with ProteinMPNN(*24*) on a fixed backbone, and mutation-induced changes were reported as ΔlogP relative to WT. Thermodynamic stability changes (ΔΔG) were predicted using ThermoMPNN(*25*) under an identical fixed-backbone assumption, with positive values indicating destabilisation. To ensure robustness, the full analysis was independently repeated using alternative inverse folding and stability prediction frameworks, including ESM-IF(*31*) and FoldX(*47*). In addition to AlphaMissense(*29*), mutation effect scores derived from ESM-1b(*33, 48*) were analysed and are reported in the Supplementary Information. Across all methods, the qualitative structure of the ΔlogP-ΔΔG landscape and the enrichment patterns of pathogenic mutations were conserved.

Predicted pathogenicity scores were mapped directly onto the ΔlogP-ΔΔG landscape. Rather than imposing arbitrary energetic thresholds, a data-driven reparameterisation was applied by translating each axis to the intersection point of the pathogenic and benign score distributions, preserving the physical interpretation of both metrics while capturing IP_3_R1-specific biophysical constraint structure.

### Ensemble generation, pocket definition, and conformational substate analysis

Conformational ensembles of the R1-NT were generated using BioEmu. For each system, including WT and selected disease-associated variants, 1,000 conformations were sampled under identical conditions. Side chains were reconstructed for all generated conformations prior to analysis. To ensure the sampling of physiologically relevant ensembles, inter-domain contacts between the suppressor domain (SD, residues 1-225) and the IP_3_-binding core (IBC, residues 226-604) were quantified using a heavy-atom distance cutoff. Conformations were restricted to topologically native states by excluding outlier frames where inter-domain contact densities fell below a statistically defined threshold. This filtering step effectively removed dissociated or non-native configurations that deviate from the canonical receptor architecture, thereby preserving the structural integrity of the functional module for subsequent analysis.

Subsequent analyses focused on the IP_3_-binding pocket, defined by residues 265, 267-269, 508, 511, and 567-569. Each retained conformation was represented using a reduced pocket-local feature set comprising backbone Cartesian coordinates, backbone dihedral angles (𝜑, 𝜓), and side-chain 𝜒_!_rotamers of pocket-forming residues. Low-dimensional embeddings were constructed using UMAP(*36*), followed by density-based clustering using DBSCAN(*37*) to identify discrete pocket conformational substates without pre-specifying cluster number. Noise points were excluded, and across parameter choices three reproducible, well-separated pocket substates were consistently identified.

To ensure structural equivalence of the binding site across systems, ligand-binding pockets were identified on a frame-by-frame basis using fpocket(*40*), retaining only pockets fully matching the predefined residue set. Quantitative pocket descriptors, including pocket volume, solvent-accessible surface area, hydrophobicity, and fpocket score, were extracted for all retained conformations. For each system, the fractional occupancy of each pocket substate was computed and compared to the WT ensemble to quantify mutation-induced redistribution of conformational populations. Differences in pocket property distributions between WT and mutants were assessed using non-parametric Mann-Whitney U tests with false-discovery-rate correction.

### Molecular dynamics simulations and adaptive sampling

Representative conformations for molecular dynamics simulations were selected from the centres of the BioEmu-derived pocket substates for each system, ensuring structurally comparable starting points across variants. All-atom molecular dynamics simulations were performed using the ACEMD engine(*49*) with the CHARMM36m force field(*50*) and the TIP3P water model(*51*). Each system was solvated in an explicit periodic box of water with a minimum padding of 10 Å from any protein atom, neutralised, and ionised to a final NaCl concentration of 0.15 M.

Energy minimisation was carried out for 5,000 steps to remove local steric clashes, followed by restrained equilibration under NPT conditions at 300 K for 5 ns. Positional restraints were applied to protein Cα atoms and side-chain heavy atoms during equilibration and subsequently released. Production simulations were performed in the NVT ensemble using a Langevin thermostat with a damping coefficient of 0.1 ps^-1^. Hydrogen mass repartitioning was employed to enable a 4-fs integration timestep(*52*). Long-range electrostatic interactions were computed using the particle-mesh Ewald(*53*) method, and periodic boundary conditions were applied in all directions.

Adaptive molecular dynamics simulations were conducted using an MSM-based adaptive sampling framework implemented with ACEMD(*49, 54, 55*). Initial seeds were propagated in multiple rounds of short simulations, with new simulations initiated from under-sampled regions of conformational space. Adaptive sampling was guided by pocket-local collective variables defined as explicit pairwise Cα-Cα distances among pocket-forming residues. For each system, adaptive sampling proceeded until an aggregate simulation time of at least 2 μs was reached, ensuring sufficient exploration of the pocket free energy landscape.

### Markov state modelling and kinetic analysis

Markov state models (MSMs) were constructed using PyEMMA (version 2.5.12)(*42*) to characterise the kinetically relevant conformational states and their interconversion from the adaptive molecular dynamics trajectories. To capture the slow dynamical modes of the IP_3_-binding pocket, each trajectory frame was represented using a pocket-local feature set comprising explicit pairwise Cα-Cα distances among pocket-forming residues and side-chain 𝜒_!_dihedral angles of the same residues. Feature selection was guided by variational approach for Markov processes (VAMP-2) scores, and the selected feature set consistently maximised kinetic variance across systems.

The resulting high-dimensional feature space was projected onto a low-dimensional kinetic subspace using time-lagged independent component analysis (TICA)(*43*). Conformations were discretised into microstates by k-means clustering in the TICA space, such that conformations within the same microstate were geometrically similar and rapidly interconverting. Transition probability matrices between microstates were estimated using a Bayesian framework at an appropriate lag time(*44*). The lag time was selected based on convergence of implied timescales, indicating that the dynamics had become approximately Markovian.

Microstates were further grouped into a small number of metastable macrostates using Perron cluster cluster analysis (PCCA), based on kinetic similarity. Model validity was assessed using Chapman-Kolmogorov tests, confirming agreement between MSM-predicted and directly observed transition probabilities over longer timescales.

Equilibrium free energies of metastable states were computed from the stationary MSM probabilities. For each macrostate 𝑆_*i*_, the free energy was calculated as

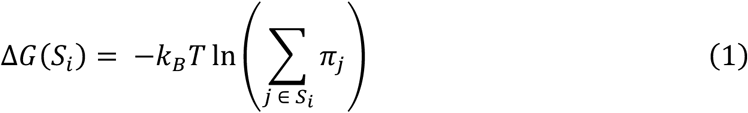

where 𝜋*_j_* denotes the stationary probability of microstate 𝑗, 𝑘*_B_* is the Boltzmann constant, and 𝑇 is the temperature. Mean first-passage times (MFPTs) between macrostates were computed using the Bayesian MSM framework, providing quantitative measures of mutation-induced changes in kinetic connectivity and conformational exchange. MSM-derived metastable states were interpreted as kinetically coherent ensembles and were not assumed to correspond directly to the geometrically defined pocket substates identified in BioEmu analyses.

### Dynamical network construction and allosteric communication analysis

Residue-level dynamical networks were constructed from molecular dynamics trajectories to quantify long-range allosteric communication between the SD and the IBC. Network nodes corresponded to protein residues, and edges were defined by persistent Cα-Cα contacts using a distance cutoff of 4.5 Å. Only contacts present for a substantial fraction of the trajectory were retained, ensuring that edges reflected physically meaningful and recurrent interactions rather than transient encounters.

To incorporate dynamical coupling information, dynamic cross-correlation matrices (DCCMs) were computed from the adaptive molecular dynamics trajectories using the Bio3D package(*56*). Correlations were calculated from atomic fluctuations about mean positions after removal of overall translation and rotation, yielding a normalised cross-correlation coefficient 𝐶*_ij_* for each residue pair 𝑖 and 𝑗. This approach provides a physically grounded measure of concerted motion derived directly from the MD trajectories(*57*).

Dynamical information was integrated into the residue interaction networks by weighting each persistent contact using the magnitude of the corresponding DCCM element. Specifically, edge weights were defined as

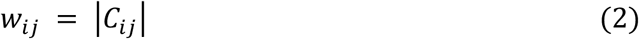

such that residue pairs exhibiting strongly correlated or anti-correlated motions contributed more strongly to communication, independent of correlation sign.

To enable path-based and flow-based analyses, correlation-weighted networks were converted into effective communication graphs by defining edge lengths as the negative logarithm of the weights,

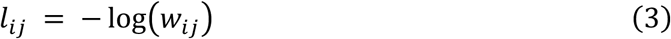

which maps strongly coupled interactions to short effective distances and weakly coupled interactions to longer distances. In this formulation, signal propagation between distant regions can be interpreted in terms of cumulative communication cost along network paths.

Allosteric communication between defined source residues in the SD and target residues forming the IP_3_-binding pocket was quantified using current-flow (random-walk)(*58*) betweenness centrality. In this framework, the residue interaction network is treated as an electrical resistor network in which edges conduct current inversely proportional to their effective lengths. The current-flow betweenness(*59*) of a node or edge corresponds to the expected amount of current passing through it when unit current is injected at the source residues and removed at the target residues, averaged over all possible paths connecting the two regions. Unlike shortest-path approaches, this metric integrates contributions from the entire ensemble of communication pathways and is therefore sensitive to global redistribution of signalling rather than reliance on a single optimal route.

To assess robustness and temporal variability, trajectories were subdivided into multiple time blocks and analysed independently. Residue- and edge-level current-flow betweenness values were computed for each block and averaged across blocks. In parallel, the shortest effective path length between source and target regions was computed as

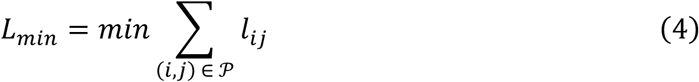

where 𝒫 denotes all possible paths connecting source and target residues. Comparisons between WT and mutant systems were performed across blocks to identify systematic mutation-induced redistribution of allosteric communication pathways and changes in overall signalling efficiency.

### Computational analysis and visualisation

All computational analyses were performed using in-house Python scripts with standard scientific libraries. Molecular dynamics trajectories were processed and analysed using MDAnalysis(*60*) and MDTraj(*61*). These tools were used to compute structural and dynamical observables, including root-mean-square deviation (RMSD), root-mean-square fluctuation (RMSF), radius of gyration (Rg), solvent-accessible surface area (SASA), residue-residue contact patterns, backbone dihedral angles (𝜑, 𝜓), and side-chain 𝜒_1_dihedral angles. These metrics were used to quantify both global structural stability and pocket-local conformational variability across systems.

Numerical analysis, data handling, and statistical evaluation were performed using NumPy(*62*), SciPy(*63*), Pandas(*64*), and scikit-learn(*65*). Figures were generated using Matplotlib(*66*), with consistent styling applied across datasets to facilitate direct comparison between conditions. Molecular structures, conformational ensembles, and representative trajectories were visualised using UCSF ChimeraX(*14*) and Protein Imager(*67*). These tools were used to inspect pocket geometries, domain interfaces, and mutation-induced structural changes, and to generate molecular renderings used in figures.

## Acknowledgements

The authors gratefully acknowledge support from the David James Fund, Department of Pharmacology. Y.Z. also acknowledges personal financial support from the Percy Lander Studentship, Downing College, Cambridge.

