## Supplemental Materials for "A unified ensemble-allosteric framework reconciles gain- and loss-of-function disease mutations in the IP_3_ receptor"

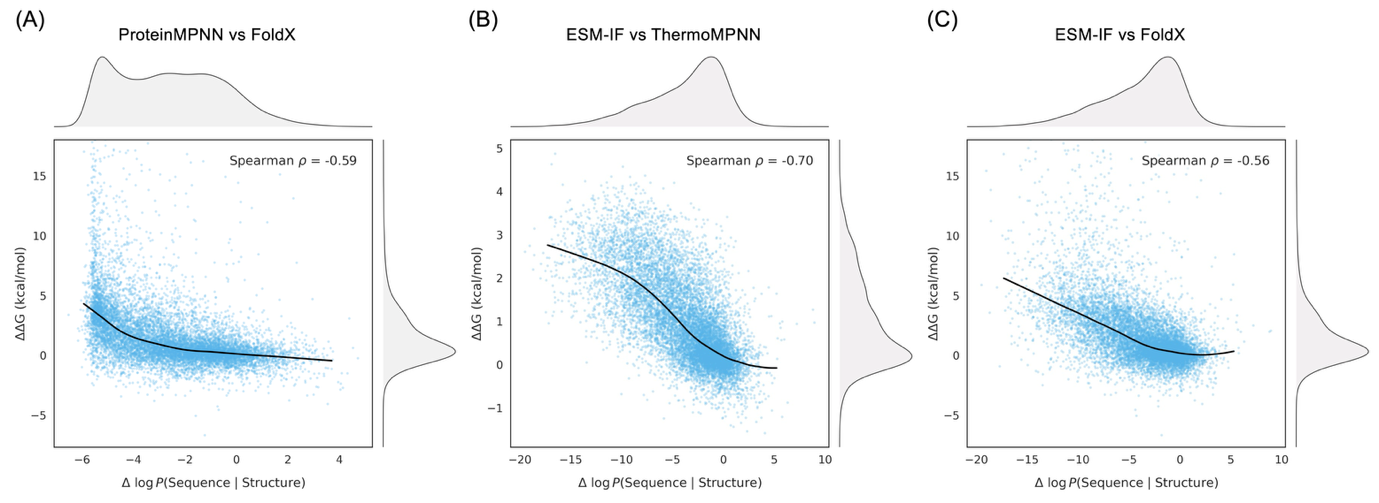

**Figure S1. Correlation of sequence-structure compatibility and thermodynamic stability across diverse model pairings.** **(A-C)** Joint plots displaying the relationship between the log-likelihood of sequence given structure (Δ log P) and change in Gibbs free energy (ΔΔG) for all possible single-point mutations in the IP_3_R1 N-terminal region (n = 604 ×19 variants). Comparisons are shown for ProteinMPNN versus FoldX (A), ESM-IF versus ThermoMPNN (B), and ESM-IF versus FoldX (C). Marginal density plots indicate the univariate distribution of each metric. Black solid lines represent a non-linear fit to the data, highlighting the general trend where mutations with lower sequence compatibility tend to exhibit higher thermodynamic destabilisation. Spearman’s ρ values are provided for each pairing, showing moderate to strong negative correlations across all tested model combinations.

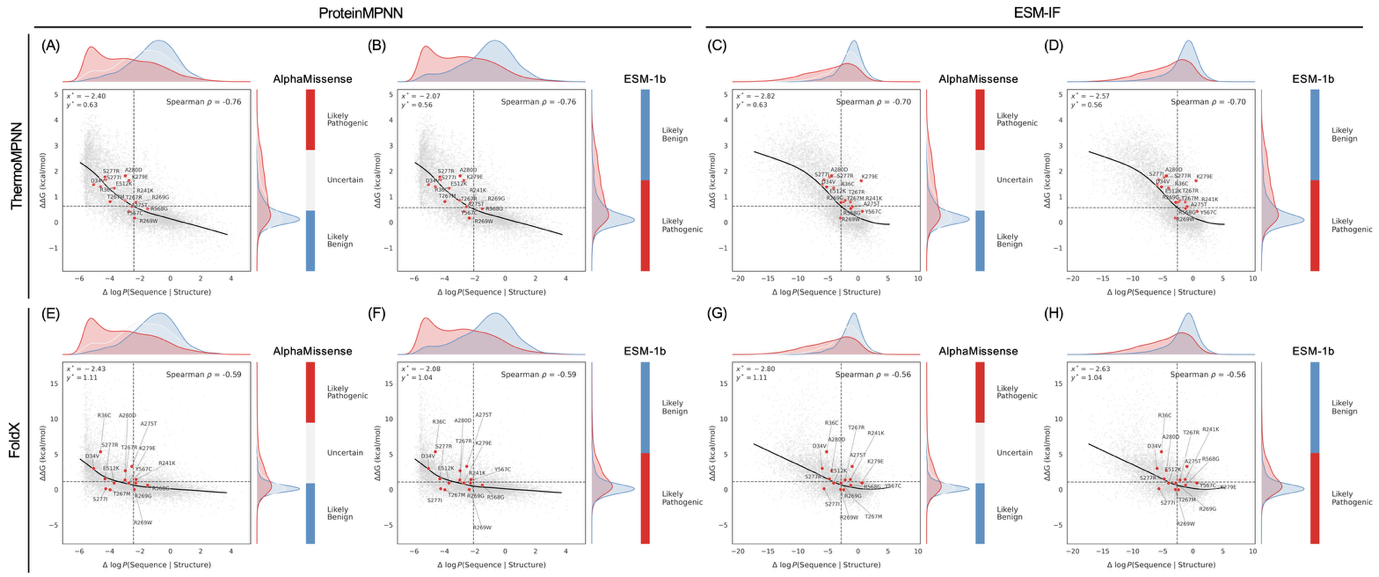

**Figure S2. Robustness of the IP_3_R1 mutational landscape across diverse computational architectures.** **(A-D)** Stability changes (ΔΔG) calculated via ThermoMPNN versus sequence-structure compatibility (Δ log P) derived from ProteinMPNN (A, B) and ESM-IF (C, D). **(E-H)** Corresponding analysis using FoldX for (ΔΔG) calculations with ProteinMPNN (E, F) and ESM-IF (G, H) likelihoods. In all panels, individual points are coloured according to their respective AlphaMissense (A, C, E, G) or ESM-1b (B, D, F, H) scores, with marginal densities showing the distribution of likely pathogenic (red) and likely benign (blue) variants. Dashed lines indicate the data-driven reference points (x*, y*) at which the clinical score distributions intersect. Labelled dots denote known gain- and loss-of-function variants. The consistent clustering of these variants across all model combinations demonstrates that the identified mutational regime is a robust structural property of the IP_3_R1 N-terminal region and not an artefact of specific predictive frameworks.

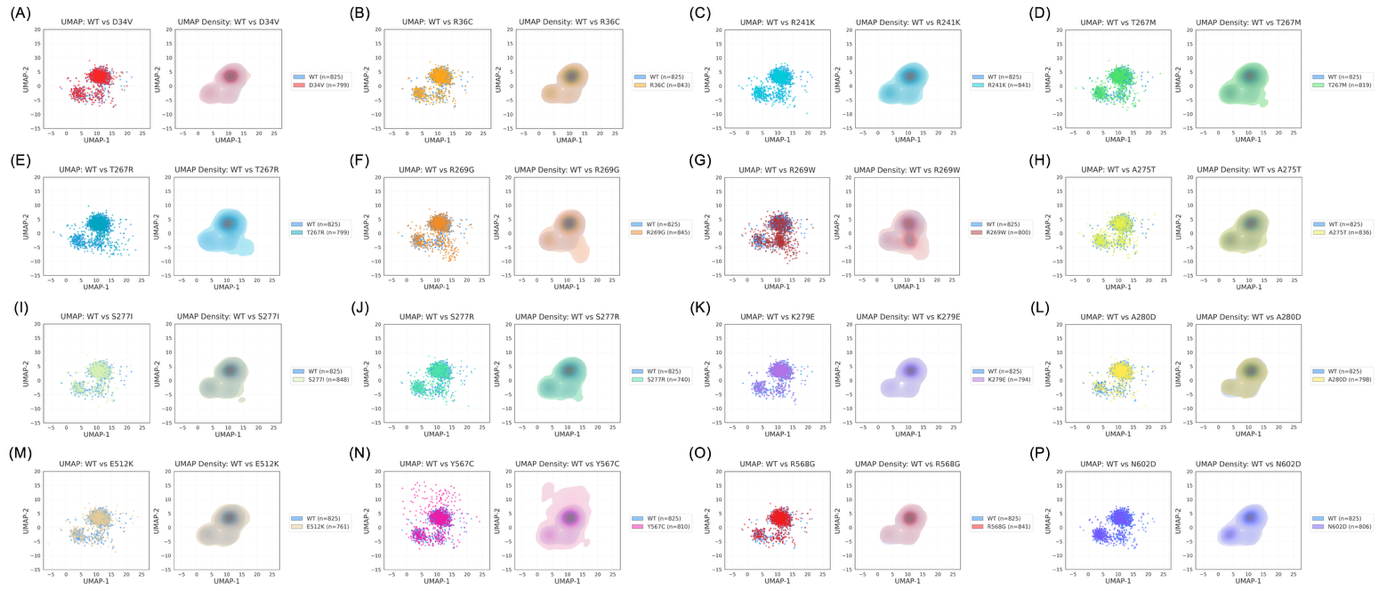

**Figure S3. Conformational distributions of IP_3_R1 NT variants.** **(A-P)** UMAP scatter plots (left) and corresponding kernel density estimates (right) comparing conformational ensembles of wild type (WT; blue) to 16 disease-associated variants. Each point represents an individual sampled conformation. The number of conformations retained post-filtering varies by system and is indicated in each respective legend. Mutants are coloured individually: **(A)** D34V; **(B)** R36C; **(C)** R241K; **(D)** T267M; **(E)** T267R; **(F)** R269G; **(G)** R269W; **(H)** A275T; **(I)** S277I; **(J)** S277R; **(K)** K279E; **(L)** A280D; **(M)** E512K; **(N)** Y567C; **(O)** R568G; **(P)** N602D.

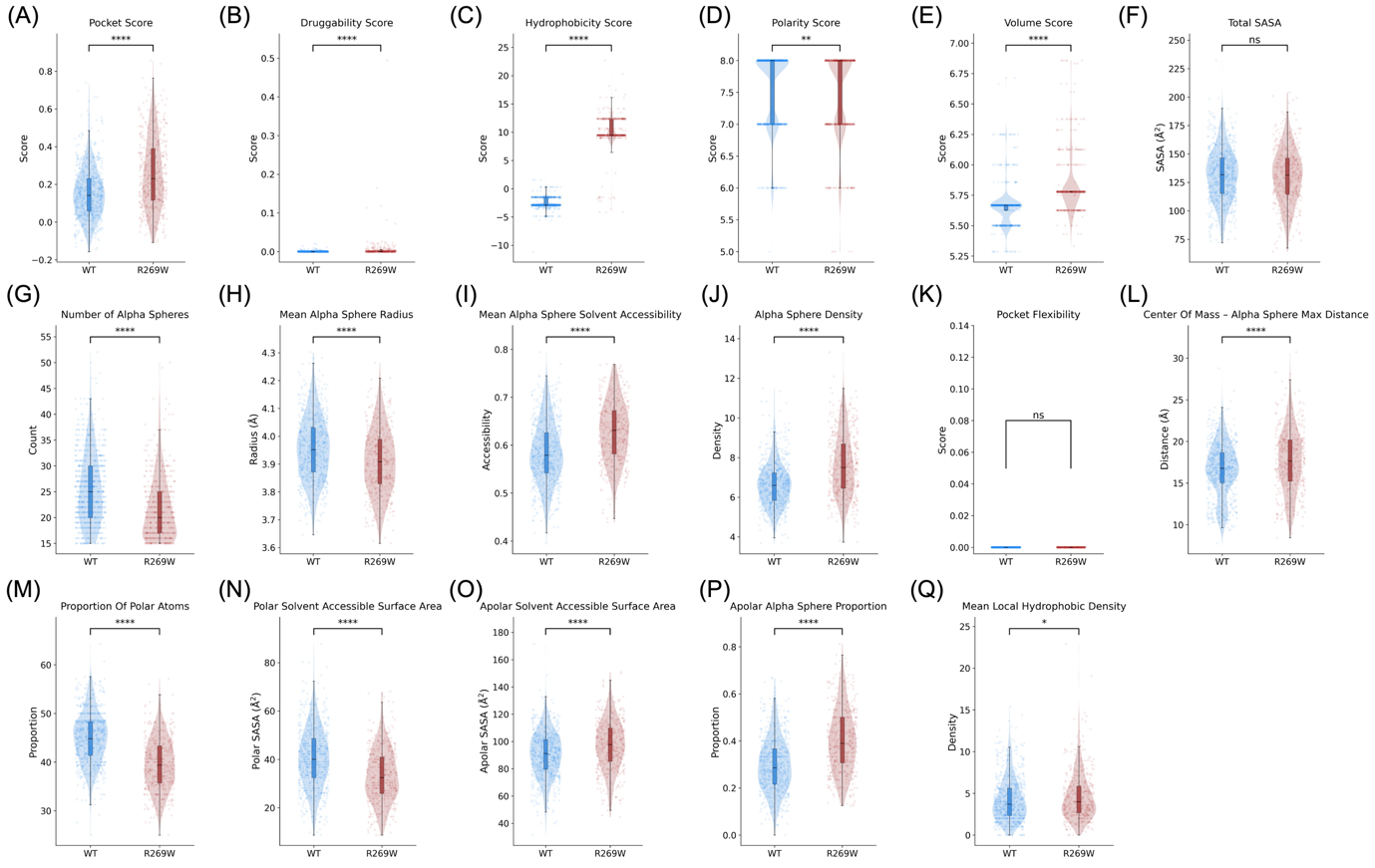

**Figure S4. Comparative physicochemical and structural descriptors of the WT and R269W binding pockets.** **(A-Q)** Violin plots illustrating the distributions of fpocket-derived parameters calculated for the wild type (WT; blue) and the R269W variant (red) conformational ensembles. Overlaid scatter points represent individual sampled conformations. Evaluated metrics include: **(A)** Pocket Score; **(B)** Druggability Score; **(C)** Hydrophobicity Score; **(D)** Polarity Score; **(E)** Volume Score; **(F)** Total solvent-accessible surface area (SASA); **(G)** Number of Alpha Spheres; **(H)** Mean Alpha Sphere Radius; **(I)** Mean Alpha Sphere Solvent Accessibility; **(J)** Alpha Sphere Density; **(K)** Pocket Flexibility; **(L)** Center of Mass to Alpha Sphere Max Distance; **(M)** Proportion of Polar Atoms; **(N)** Polar SASA; **(O)** Apolar SASA; **(P)** Apolar Alpha Sphere Proportion; and **(Q)** Mean Local Hydrophobic Density. Statistical significance was assessed using a Mann-Whitney U test (ns, not significant; *P < 0.05; **P < 0.01; ****P < 0.0001).

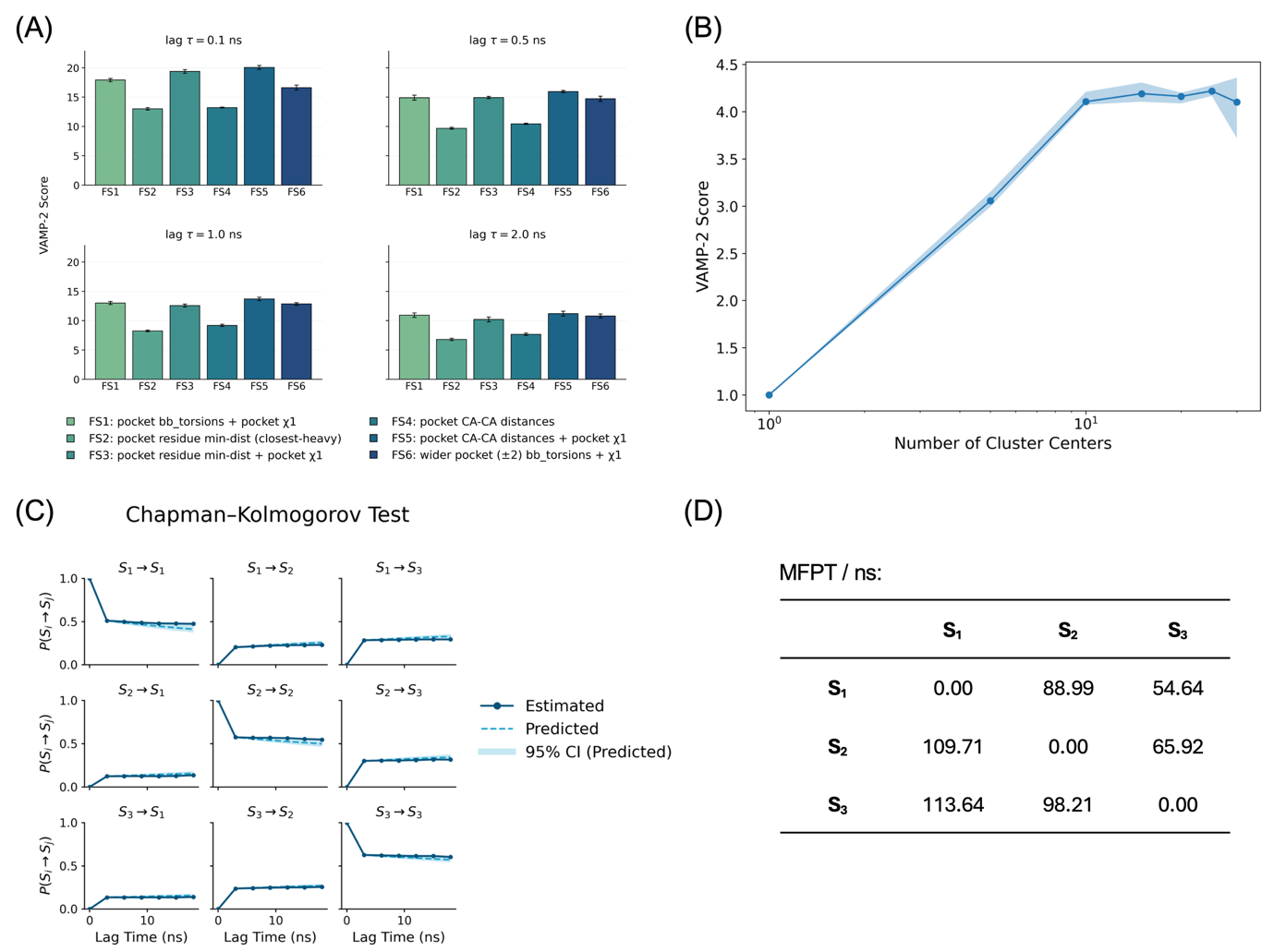

**Figure S5. Feature selection, discretisation, and validation of the wild-type Markov state model.** **(A)** VAMP-2 scores evaluating alternative pocket-local feature sets across multiple lag times. The assessed sets encompass backbone torsions, $\chi_{1}$ rotamers, minimum residue distances, Cα-Cα distances, and wider-pocket descriptors. **(B)** VAMP-2 score dependence on the number of cluster centres used for spatial discretisation. **(C)** Chapman-Kolmogorov validation of the selected three-state model, comparing empirical estimates with model-predicted transition probabilities across varying lag times. **(D)** Pairwise mean first-passage time (MFPT) matrix for the wild-type metastable states.

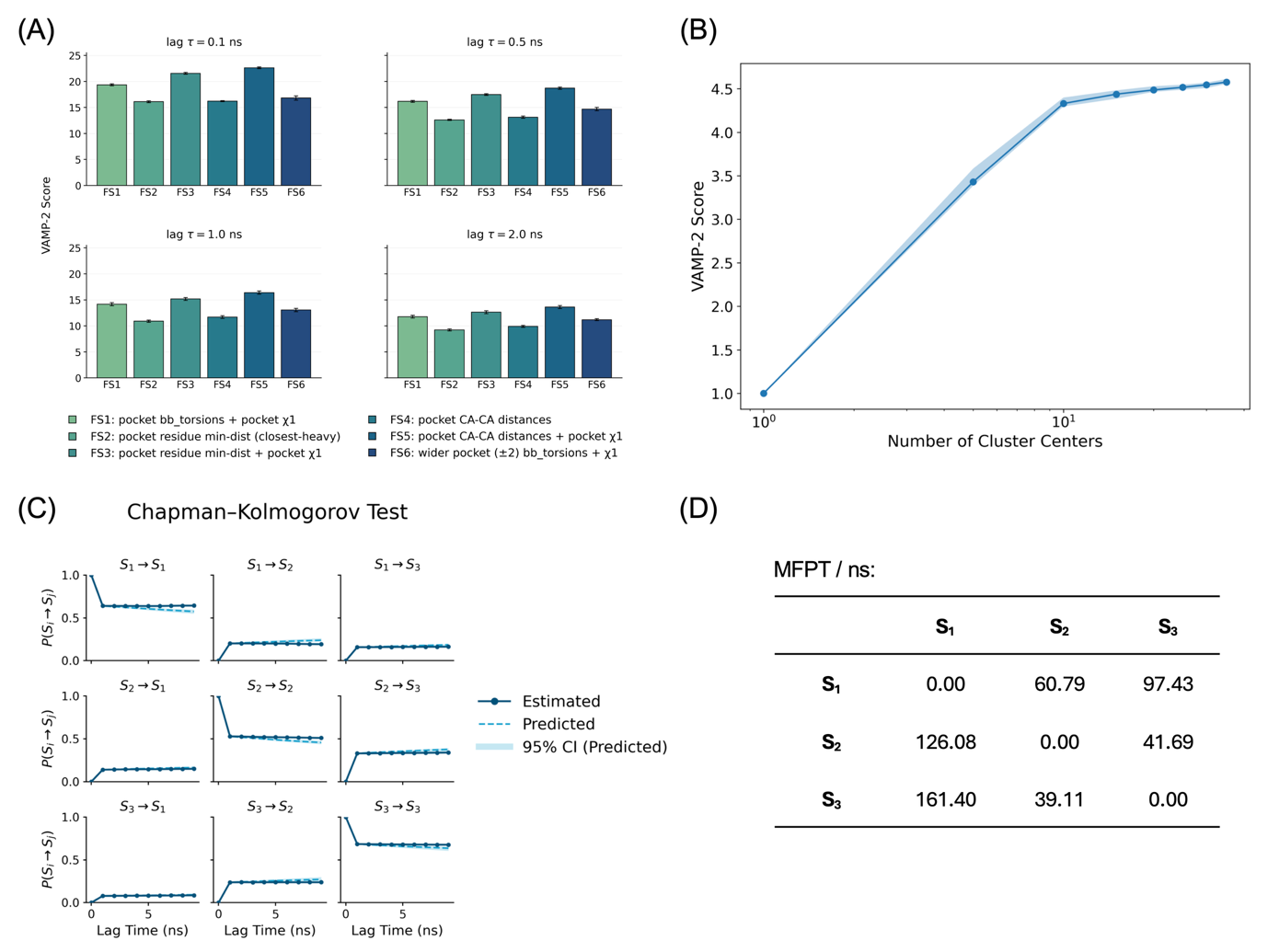

**Figure S6. Feature selection, discretisation, and validation of the R269W Markov state model. (A)** VAMP-2 scores evaluating alternative pocket-local feature sets across multiple lag times. **(B)** VAMP-2 score dependence on the number of cluster centres used for spatial discretisation. **(C)** Chapman-Kolmogorov validation of the selected three-state R269W model, comparing empirical estimates with model-predicted transition probabilities. **(D)** Pairwise mean first-passage time (MFPT) matrix for the assigned R269W metastable states.

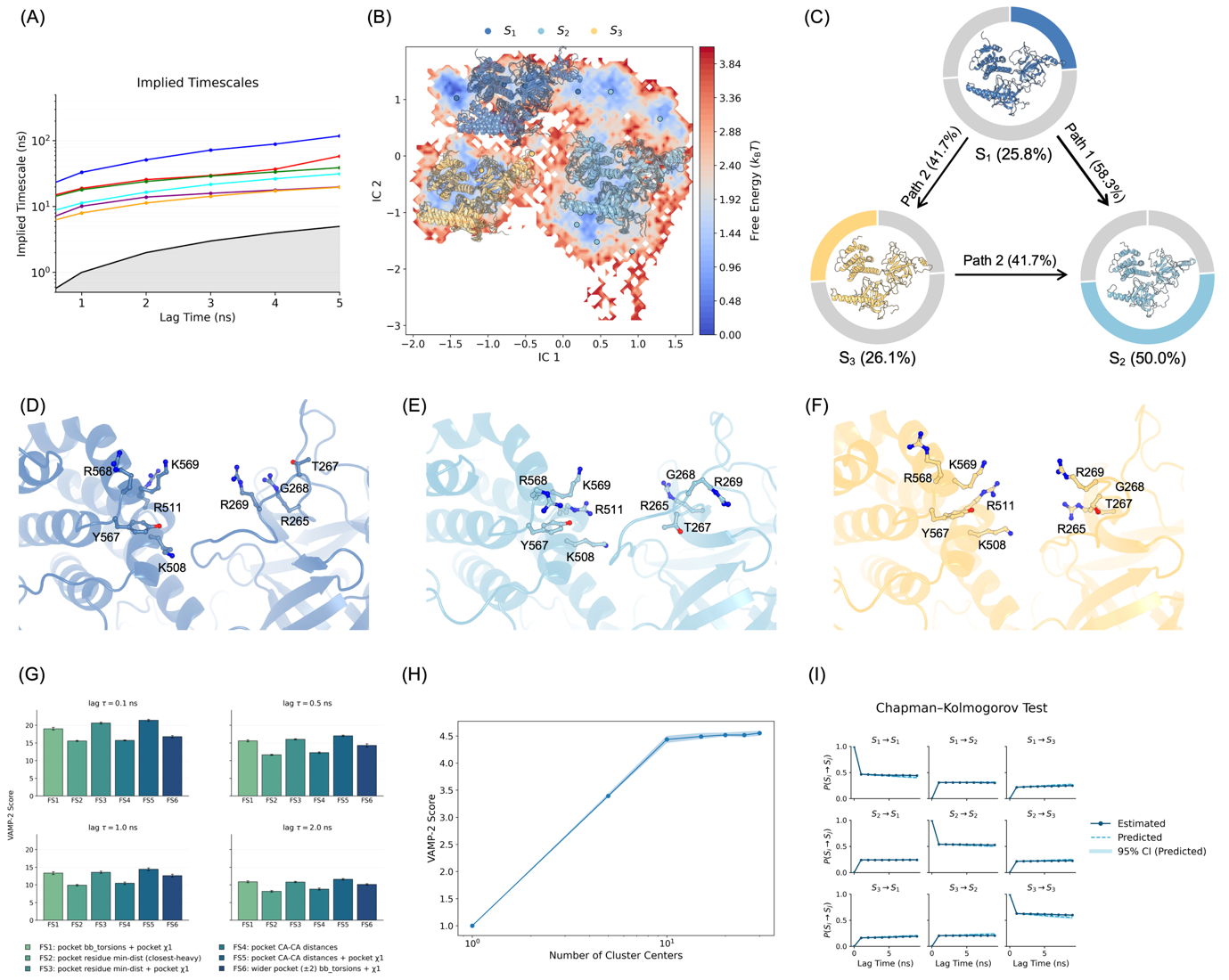

**Figure S7. Local pocket kinetics remain comparatively accessible in the R36C variant.** **(A)** Implied timescale analysis of the R36C Markov state model (MSM). **(B)** Projection of R36C adaptive sampling trajectories onto the first two independent components (tICs), overlaid with metastable state assignments. **(C)** Transition path decomposition from the least- to the most-populated state. The flux network reveals predominantly direct exchange, with a modest increase in intermediate-mediated contributions relative to the wild type. **(D-F)** Representative conformations of the R36C metastable states, focusing on the predefined IP_3_-binding pocket. **(G-I)** Feature selection (G), spatial discretisation (H), and Chapman-Kolmogorov validation (I) for the R36C model. These analyses confirm that R36C alters local kinetic exchange more subtly than R269W, avoiding the formation of a strongly rerouted or occluded pocket state.

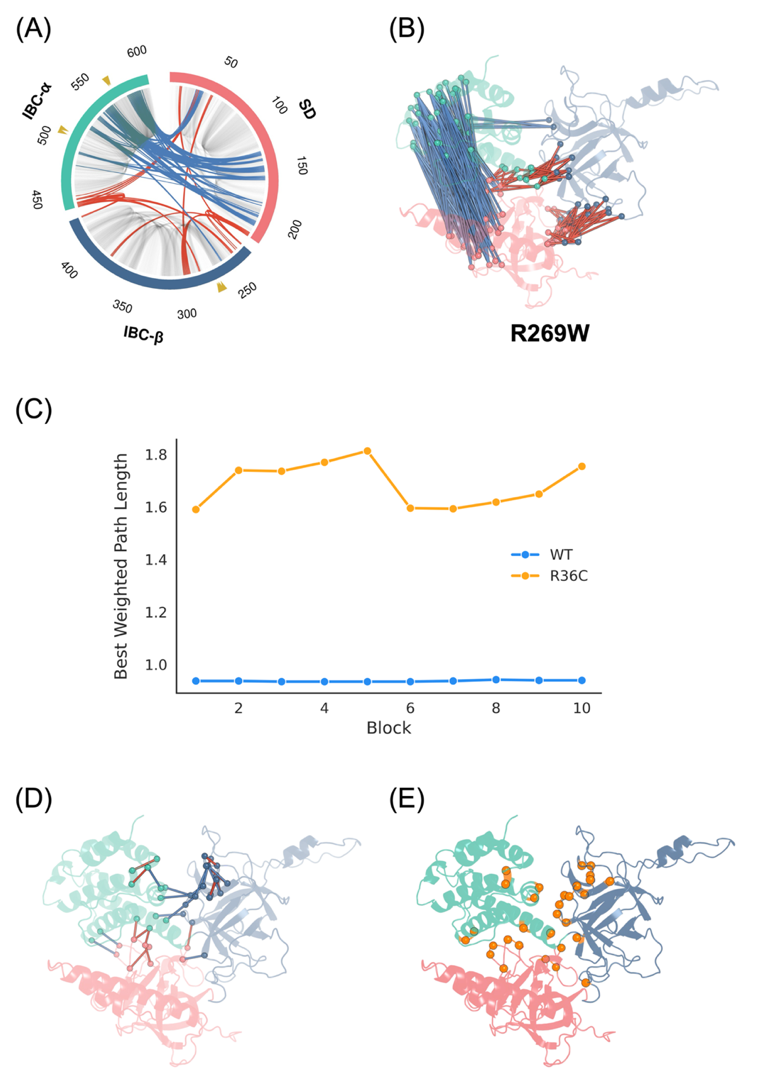

**Figure S8. The R269W mutation exhibits extensive correlation remodelling, whereas R36C increases source-to-pocket communication costs without global dynamic disruption. (A)** Domain-resolved dynamic cross-correlation analysis of the R269W mutant, illustrating extensive reorganisation of correlated motions across the SD and the IBC. **(B)** Structural mapping of prominent alterations in the R269W correlation network, projected onto the NT. **(C)** Block-resolved optimal weighted source-to-pocket pathway lengths for the wild-type (WT) and R36C. The R36C variant exhibits consistently extended pathways across trajectory blocks, indicative of attenuated communication efficiency. **(D, E)** Structural mapping of residues and inter-residue contacts demonstrating altered current-flow betweenness contributions in the R36C mutant relative to the WT.

**Table S1. Fractional occupancy of pocket conformational substates across wild type and mutant variants.**

| System | Cluster A | Cluster B | Cluster C |
| --- | --- | --- | --- |
| WT | 0.875 | 0.106 | 0.019 |
| D34V | 0.866 | 0.113 | 0.021 |
| R36C | 0.859 | 0.123 | 0.018 |
| R241K | 0.830 | 0.148 | 0.022 |
| T267R | 0.890 | 0.083 | 0.027 |
| T267M | 0.818 | 0.168 | 0.014 |
| R269G | 0.855 | 0.118 | 0.028 |
| R269W | 0.749 | 0.128 | 0.123 |
| A275T | 0.784 | 0.185 | 0.031 |
| S277I | 0.852 | 0.128 | 0.020 |
| S277R | 0.845 | 0.140 | 0.015 |
| K279E | 0.883 | 0.096 | 0.021 |
| A280D | 0.914 | 0.069 | 0.017 |
| E512K | 0.878 | 0.104 | 0.018 |
| Y567C | 0.859 | 0.116 | 0.025 |
| R568G | 0.918 | 0.062 | 0.019 |
| N602D | 0.717 | 0.264 | 0.020 |

**Table S2. Comparative physicochemical analysis of the IP_3_ binding pocket in WT and R269W variants.**

| Feature | WT  Median | R269W Median | Δ Median | Cliff's Delta (δ) | p value | FDR |
| --- | --- | --- | --- | --- | --- | --- |
| Charge score | 6.000 | 5.000 | -1.000 | 1.000 | 5.48×10^-156^ | 3.47×10^-155^ |
| Volume score | 5.667 | 5.778 | 0.111 | -1.000 | 5.48×10^-156^ | 3.47×10^-155^ |
| Hydrophobicity score | -2.889 | 9.444 | 12.333 | -1.000 | 5.48×10^-156^ | 3.47×10^-155^ |
| Proportion of polar atoms | 45.455 | 40.625 | -4.830 | 0.544 | 1.36×10^-33^ | 6.45×10^-33^ |
| Alpha sphere density | 6.639 | 7.544 | 0.905 | -0.414 | 3.65×10^-20^ | 1.35×10^-19^ |
| Apolar alpha sphere proportion | 0.284 | 0.375 | 0.091 | -0.414 | 4.28×10^-20^ | 1.35×10^-19^ |
| Number of Alpha Spheres | 27.000 | 21.000 | -6.000 | 0.367 | 3.90×10^-16^ | 1.06×10^-15^ |
| Mean alp. sph. solvent access | 0.578 | 0.614 | 0.037 | -0.326 | 4.58×10^-13^ | 1.09×10^-12^ |
| Apolar SASA | 92.987 | 102.648 | 9.661 | -0.323 | 7.65×10^-13^ | 1.62×10^-12^ |
| Druggability Score | 0.000 | 0.000 | 0.000 | -0.221 | 7.53×10^-11^ | 1.43×10^-10^ |
| Polar SASA | 42.130 | 35.646 | -6.484 | 0.283 | 3.18×10^-10^ | 5.49×10^-10^ |
| Volume | 677.265 | 607.406 | -69.859 | 0.278 | 7.23×10^-10^ | 1.15×10^-9^ |
| Score | 0.146 | 0.217 | 0.071 | -0.275 | 1.01×10^-9^ | 1.48×10^-9^ |
| Mean alpha sphere radius | 3.961 | 3.914 | -0.046 | 0.198 | 1.13×10^-5^ | 1.53×10^-5^ |
| Cent. of mass - Alpha Sphere max dist | 16.860 | 17.505 | 0.645 | -0.134 | 0.003 | 0.004 |
| Total SASA | 134.567 | 137.745 | 3.178 | -0.087 | 0.054 | 0.065 |
| Mean local hydrophobic density | 3.750 | 3.800 | 0.050 | -0.021 | 0.647 | 0.723 |
| Polarity score | 8.000 | 8.000 | 0.000 | 0.000 | 1 | 1 |
| Flexibility | 0.000 | 0.000 | 0.000 | 0.000 | 1 | 1 |

**Table S3. MSM thermodynamic and kinetic parameters for WT, R269W and R36C.**

**Table S3A. State populations and state-to-complement kinetics.**

| System | State | Population | Relative Free Energy (G/kT) | State-to-complement MFPT (ns) | Complement-to-state MFPT (ns) |
| --- | --- | --- | --- | --- | --- |
| WT | S_1_ | 0.258 | 1.353 | 33.5 ± 7.5 | 121.2 ± 22.1 |
| WT | S_2_ | 0.259 | 1.350 | 45.0 ± 8.5 | 105.4 ± 19.2 |
| WT | S_3_ | 0.483 | 0.729 | 122.1 ± 22.8 | 57.6 ± 16.2 |
| R269W | S_1_ | 0.288 | 1.245 | 58.6 ± 9.6 | 152.1 ± 26.7 |
| R269W | S_2_ | 0.299 | 1.206 | 17.2 ± 1.6 | 48.8 ± 6.1 |
| R269W | S_3_ | 0.413 | 0.885 | 166.8 ± 27.1 | 99.5 ± 13.2 |
| R36C | S_1_ | 0.239 | 1.433 | 25.2 ± 3.5 | 88.6 ± 16.2 |
| R36C | S_2_ | 0.501 | 0.691 | 54.2 ± 10.1 | 55.4 ± 9.8 |
| R36C | S_3_ | 0.261 | 1.345 | 72.5 ± 13.4 | 55.3 ± 8.5 |

**Table S3B. Transition-path decomposition from least-populated state to dominant state.**

| System | Least-populated State | Dominant State | Direct Path Flux | Intermediate-mediated Path Flux |
| --- | --- | --- | --- | --- |
| WT | S_1_ | S_3_ | 80.0% | 20.0% |
| R269W | S_1_ | S_3_ | 13.2% | 86.8% |
| R36C | S_1_ | S_2_ | 58.3% | 41.7% |

**Table S3C. Pairwise MFPT matrices of the three systems.**

**WT**

| From \ To | S_1_ | S_2_ | S_3_ |
| --- | --- | --- | --- |
| S_1_ | 0.00 | 88.99 | 54.64 |
| S_2_ | 109.71 | 0.00 | 65.92 |
| S_3_ | 113.64 | 98.21 | 0.00 |

**R269W**

| From \ To | S_1_ | S_2_ | S_3_ |
| --- | --- | --- | --- |
| S_1_ | 0.00 | 60.79 | 97.43 |
| S_2_ | 126.08 | 0.00 | 41.69 |
| S_3_ | 161.40 | 39.11 | 0.00 |

**R36C**

| From \ To | S_1_ | S_2_ | S_3_ |
| --- | --- | --- | --- |
| S_1_ | 0.00 | 54.42 | 52.48 |
| S_2_ | 86.86 | 0.00 | 73.40 |
| S_3_ | 66.31 | 52.20 | 0.00 |

**Table S4. Community membership of WT and R36C contact-informed dynamical networks.**

| System | Community | Number of Residues | Functional annotation | Pocket Residues | Residue List |
| --- | --- | --- | --- | --- | --- |
| WT | C1 | 30 | IBC-α / pocket-side module | K508, R511, Y567, R568, K569 | V481, V484, P502, R504, E505, R506, Q507, K508, L509, M510, R511, E512, Q513, I515, L516, L560, R561, H562, S563, Q564, Q565, D566, Y567, R568, K569, N570, Q571, E572, I574, L598 |
| WT | C2 | 16 | SD-IBC interface / distal IBC-β | — | L57, K127, V221, L222, F223, M224, K235, V284, E285, V287, C292, R293, G294, G295, G297, V433 |
| WT | C3 | 24 | SD community | — | A19, N24, I27, S28, T29, L30, G31, V33, D34, D35, R36, C37, V38, V39, Q40, D55, V149, T150, L151, D152, A195, C206, E208, V209 |
| WT | C4 | 19 | IBC-β pocket loop | R265, T267, G268, R269 | A245, K249, L251, T252, F263, L264, R265, T266, T267, G268, R269, Q270, S271, A272, T273, L355, S401, M415, L416 |
| WT | C5 | 16 | IBC-α interdomain bridge | — | L32, S128, A438, E439, R441, D442, L443, D444, F445, A446, N447, D448, S450, K451, V452, N514 |
| WT | C6 | 13 | IBC-β pocket-adjacent module | — | R241, L242, F250, S274, A275, T276, S277, S278, K279, A280, L281, H307, A309 |
| WT | C7 | 9 | IBC-α peripheral module | — | A449, L453, L476, L477, E478, D479, L480, I519, V559 |
| WT | C8 | 7 | SD-IBC bridge | — | K52, C56, G237, V239, E283, L308, V440 |
| R36C | C1 | 26 | IBC-α / pocket-side module | K508, R511, Y567, R568, K569 | V481, V484, R504, E505, Q507, K508, L509, M510, R511, E512, Q513, I515, L516, L560, R561, H562, S563, Q564, D566, Y567, R568, K569, N570, E572, I574, L598 |
| R36C | C2 | 16 | SD-IBC interface / distal IBC-β | — | L57, K127, V221, L222, F223, M224, K235, G237, V284, E285, V287, C292, R293, G294, G295, G297 |
| R36C | C3 | 23 | SD community | — | I27, S28, T29, L30, G31, L32, V33, D34, D35, C36, C37, V38, Q40, K52, D55, C56, S128, V149, T150, L151, D152, E208, V209 |
| R36C | C4 | 14 | IBC-β pocket loop | T267, G268, R269 | L264, T266, T267, G268, R269, Q270, S271, A272, T273, S274, A275, S401, M415, L416 |
| R36C | C5 | 5 | IBC-α interdomain fragment | — | D448, S450, K451, V452, N514 |
| R36C | C6 | 18 | IBC-β pocket-adjacent module | R265 | L242, A245, K249, F250, T252, F263, R265, T276, S277, S278, K279, A280, L281, E283, H307, L308, A309, L355 |
| R36C | C7 | 6 | IBC-α peripheral module | — | L476, L477, E478, D479, L480, V559 |
| R36C | C8 | 17 | IBC-α / IBC-β bridge | — | V239, R241, L251, V433, A438, E439, V440, R441, D442, L443, D444, F445, A446, N447, A449, L453, I519 |
| R36C | C9 | 5 | SD fragment | — | A19, N24, V39, A195, C206 |
| R36C | C10 | 4 | IBC-α fragment | — | P502, R506, Q565, Q571 |

**Table S5. Top residue-level current-flow changes between WT and R36C.**

| Rank | Residue | Domain | Δ current-flow (R36C − WT) | 95% CI lower | 95% CI upper |
| --- | --- | --- | --- | --- | --- |
| 1 | G311 | IBC-β | 1.75 x 10^-5^ | 1.59 x 10^-5^ | 1.91 x 10^-5^ |
| 2 | K51 | SD | 1.56 x 10^-5^ | 1.37 x 10^-5^ | 1.74 x 10^-5^ |
| 3 | S277 | IBC-β | -1.50 x 10^-5^ | -1.87 x 10^-5^ | -1.07 x 10^-5^ |
| 4 | R265 | IBC-β | 1.43 x 10^-5^ | 1.36 x 10^-5^ | 1.50 x 10^-5^ |
| 5 | A275 | IBC-β | -1.22 x 10^-5^ | -1.68 x 10^-5^ | -7.72 x 10^-6^ |
| 6 | L509 | IBC-α | -1.18 x 10^-5^ | -1.48 x 10^-5^ | -8.18 x 10^-6^ |
| 7 | L32 | SD | 1.16 x 10^-5^ | 8.12 x 10^-6^ | 1.50 x 10^-5^ |
| 8 | T276 | IBC-β | -1.14 x 10^-5^ | -1.58 x 10^-5^ | -6.51 x 10^-6^ |
| 9 | T267 | IBC-β | -9.82 x 10^-6^ | -1.15 x 10^-5^ | -8.00 x 10^-6^ |
| 10 | V33 | SD | 9.66 x 10^-6^ | 7.32 x 10^-6^ | 1.18 x 10^-5^ |
| 11 | D202 | SD | -9.64 x 10^-6^ | -1.10 x 10^-5^ | -8.21 x 10^-6^ |
| 12 | V201 | SD | -9.38 x 10^-6^ | -1.04 x 10^-5^ | -7.63 x 10^-6^ |
| 13 | S278 | IBC-β | -9.13 x 10^-6^ | -1.58 x 10^-5^ | -6.62 x 10^-7^ |
| 14 | S271 | IBC-β | 8.36 x 10^-6^ | 4.54 x 10^-6^ | 1.16 x 10^-5^ |
| 15 | E469 | IBC-α | -7.99 x 10^-6^ | -8.49 x 10^-6^ | -7.62 x 10^-6^ |
| 16 | F445 | IBC-α | 7.99 x 10^-6^ | 5.90 x 10^-6^ | 1.03 x 10^-5^ |
| 17 | G44 | SD | 7.93 x 10^-6^ | 6.61 x 10^-6^ | 9.50 x 10^-6^ |
| 18 | E512 | IBC-α | -7.90 x 10^-6^ | -1.37 x 10^-5^ | -7.20 x 10^-7^ |
| 19 | R36 | SD | 7.82 x 10^-6^ | 6.05 x 10^-6^ | 9.82 x 10^-6^ |
| 20 | T266 | IBC-β | 7.32 x 10^-6^ | 4.90 x 10^-6^ | 9.64 x 10^-6^ |
| 21 | T252 | IBC-β | 7.22 x 10^-6^ | 6.00 x 10^-6^ | 8.43 x 10^-6^ |
| 22 | A309 | IBC-β | -7.21 x 10^-6^ | -1.17 x 10^-5^ | -1.21 x 10^-6^ |
| 23 | N468 | IBC-α | -6.83 x 10^-6^ | -8.51 x 10^-6^ | -5.05 x 10^-6^ |
| 24 | V239 | IBC-β | 6.68 x 10^-6^ | 3.67 x 10^-6^ | 9.38 x 10^-6^ |
| 25 | Q564 | IBC-α | 6.51 x 10^-6^ | 1.27 x 10^-6^ | 1.10 x 10^-5^ |
| 26 | V440 | IBC-α | 6.33 x 10^-6^ | 4.18 x 10^-6^ | 8.28 x 10^-6^ |
| 27 | N229 | IBC-β | -6.30 x 10^-6^ | -7.60 x 10^-6^ | -5.11 x 10^-6^ |
| 28 | S563 | IBC-α | 6.20 x 10^-6^ | 3.42 x 10^-6^ | 8.46 x 10^-6^ |
| 29 | E248 | IBC-β | -5.98 x 10^-6^ | -7.71 x 10^-6^ | -4.18 x 10^-6^ |
| 30 | T43 | SD | 5.41 x 10^-6^ | 3.85 x 10^-6^ | 6.72 x 10^-6^ |

**Table S6. Top edge-level current-flow rewiring events between WT and R36C.**

| Rank | Edge | Domain | Δ edge current-flow (R36C − WT) | 95% CI Lower | 95% CI Upper |
| --- | --- | --- | --- | --- | --- |
| 1 | K51-G311 | SD / IBC-β | 8.89 x 10^-6^ | 7.89 x 10^-6^ | 9.77 x 10^-6^ |
| 2 | S277-L509 | IBC-β / IBC-α | -7.67 x 10^-6^ | -9.58 x 10^-6^ | -5.39 x 10^-6^ |
| 3 | T267-A275 | IBC-β | -5.78 x 10^-6^ | -7.68 x 10^-6^ | -3.59 x 10^-6^ |
| 4 | T267-T276 | IBC-β | -4.78 x 10^-6^ | -8.26 x 10^-6^ | -1.46 x 10^-6^ |
| 5 | F250-T267 | IBC-β | -4.76 x 10^-6^ | -5.62 x 10^-6^ | -3.51 x 10^-6^ |
| 6 | F250-R265 | IBC-β | 4.54 x 10^-6^ | 2.70 x 10^-6^ | 5.93 x 10^-6^ |
| 7 | S277-E512 | IBC-β / IBC-α | -4.53 x 10^-6^ | -5.57 x 10^-6^ | -3.28 x 10^-6^ |
| 8 | S563-Y567 | IBC-α | 4.37 x 10^-6^ | 2.77 x 10^-6^ | 5.56 x 10^-6^ |
| 9 | R265-T267 | IBC-β | 4.26 x 10^-6^ | 3.06 x 10^-6^ | 5.29 x 10^-6^ |
| 10 | T252-R265 | IBC-β | 4.16 x 10^-6^ | 3.63 x 10^-6^ | 4.71 x 10^-6^ |
| 11 | T29-L32 | SD | 4.15 x 10^-6^ | 2.33 x 10^-6^ | 5.80 x 10^-6^ |
| 12 | A275-S277 | IBC-β | -4.12 x 10^-6^ | -5.20 x 10^-6^ | -2.92 x 10^-6^ |
| 13 | K249-T267 | IBC-β | -4.10 x 10^-6^ | -4.87 x 10^-6^ | -3.46 x 10^-6^ |
| 14 | Q564-K569 | IBC-α | 3.94 x 10^-6^ | 1.17 x 10^-6^ | 6.48 x 10^-6^ |
| 15 | D202-E469 | SD / IBC-α | -3.82 x 10^-6^ | -4.23 x 10^-6^ | -3.27 x 10^-6^ |
| 16 | A309-N447 | IBC-β / IBC-α | -3.67 x 10^-6^ | -4.22 x 10^-6^ | -3.11 x 10^-6^ |
| 17 | T266-G268 | IBC-β | 3.56 x 10^-6^ | 2.67 x 10^-6^ | 4.33 x 10^-6^ |
| 18 | T267-R269 | IBC-β | 3.43 x 10^-6^ | 2.03 x 10^-6^ | 4.70 x 10^-6^ |
| 19 | V201-N468 | SD / IBC-α | -3.33 x 10^-6^ | -4.34 x 10^-6^ | -2.24 x 10^-6^ |
| 20 | S278-L443 | IBC-β / IBC-α | -3.31 x 10^-6^ | -4.27 x 10^-6^ | -2.07 x 10^-6^ |
| 21 | R54-G237 | SD / IBC-β | -3.05 x 10^-6^ | -4.58 x 10^-6^ | -1.46 x 10^-6^ |
| 22 | V33-F445 | SD / IBC-α | 3.05 x 10^-6^ | 2.25 x 10^-6^ | 3.96 x 10^-6^ |
| 23 | T276-S278 | IBC-β | -3.05 x 10^-6^ | -4.08 x 10^-6^ | -1.96 x 10^-6^ |
| 24 | L32-F445 | SD / IBC-α | 2.91 x 10^-6^ | 2.33 x 10^-6^ | 3.50 x 10^-6^ |
| 25 | K249-R265 | IBC-β | 2.89 x 10^-6^ | 2.37 x 10^-6^ | 3.47 x 10^-6^ |
| 26 | S271-T273 | IBC-β | 2.83 x 10^-6^ | 2.19 x 10^-6^ | 3.45 x 10^-6^ |
| 27 | V33-A449 | SD / IBC-α | 2.80 x 10^-6^ | 2.48 x 10^-6^ | 3.11 x 10^-6^ |
| 28 | G44-K51 | SD | 2.78 x 10^-6^ | 2.30 x 10^-6^ | 3.33 x 10^-6^ |
| 29 | T266-L416 | IBC-β | 2.64 x 10^-6^ | 2.33 x 10^-6^ | 2.98 x 10^-6^ |
| 30 | V33-L476 | SD / IBC-α | 2.57 x 10^-6^ | 1.92 x 10^-6^ | 3.18 x 10^-6^ |

**Table S7. Best weighted paths from residue 36 to selected IP_3_-binding pocket residues.**

| Target Pocket Residue | WT Path Length | WT Best Weighted Path | R36C Path Length | R36C Best Weighted Path | Δ Length |
| --- | --- | --- | --- | --- | --- |
| R265 | 1.573 | R36 → D34 → V33 → F445 → D444 → V440 → V435 → R241 → F250 → R265 | 2.403 | C36 → C37 → S28 → C56 → L57 → C15 → F223 → G294 → G236 → V284 → E283 → W282 → L281 → A280 → T252 → L264 → R265 | 0.830 |
| T267 | 1.998 | R36 → D34 → V33 → F445 → D444 → V440 → V435 → R241 → F250 → R265 → T266 → T267 | 2.696 | C36 → C37 → S28 → C56 → L57 → C15 → F223 → G294 → G236 → V284 → E283 → W282 → L281 → A280 → T252 → L264 → R265 → T266 → T267 | 0.699 |
| G268 | 1.962 | R36 → D34 → V33 → F445 → D444 → V440 → V435 → R241 → F250 → R265 → T266 → G268 | 2.768 | C36 → C37 → S28 → C56 → L57 → C15 → F223 → G294 → G236 → V284 → E283 → W282 → L281 → A280 → T252 → L264 → R265 → T266 → G268 | 0.806 |
| R269 | 2.242 | R36 → D34 → V33 → F445 → D444 → V440 → V435 → R241 → F250 → R265 → T266 → G268 → R269 | 2.932 | C36 → C37 → S28 → C56 → L57 → C15 → F223 → G294 → G236 → V284 → E283 → W282 → L281 → A280 → T252 → L264 → R265 → T266 → G268 → R269 | 0.690 |
| K508 | 1.321 | R36 → D34 → V33 → F445 → A446 → Q513 → E512 → R511 → K508 | 2.190 | C36 → D34 → V33 → D448 → N447 → Q513 → E512 → K508 | 0.869 |
| R511 | 1.214 | R36 → D34 → V33 → F445 → A446 → Q513 → E512 → R511 | 2.067 | C36 → D34 → V33 → D448 → N447 → Q513 → E512 → R511 | 0.853 |
| K569 | 1.410 | R36 → D34 → V33 → F445 → A446 → Q513 → E512 → R511 → N570 → K569 | 2.364 | C36 → D34 → V33 → D448 → N447 → Q513 → E512 → R511 → N570 → K569 | 0.954 |

Paths were computed from residue 36 to designated IP_3_-binding pocket residues using effective edge lengths derived from contact-informed dynamic networks. Residue nomenclature corresponds to the human IP_3_R1 N-terminal (NT) sequence. Quantitative network analysis revealed that the mean source-to-pocket pathway length increased from 1.674 in the wild-type to 2.489 in the R36C variant, representing a 1.52-fold extension in communication cost.
